# Depletion-assisted multiplexed cell-free RNA sequencing reveals distinct human and microbial signatures in plasma versus extracellular vesicles

**DOI:** 10.1101/2023.01.31.526408

**Authors:** Hongke Wang, Qing Zhan, Meng Ning, Hongjie Guo, Qian Wang, Jiuliang Zhao, Pengfei Bao, Shaozhen Xing, Shanwen Chen, Shuai Zuo, Mengtao Li, Pengyuan Wang, Zhi John Lu

## Abstract

Cell-free long RNAs in human plasma and extracellular vesicles (EVs) have shown promise as biomarkers in liquid biopsy, despite their fragmented nature. To investigate these fragmented cell-free RNAs (cfRNAs), we developed a cost-effective cfRNA sequencing method called DETECTOR-seq (<u>de</u>pletion-assisted multipl<u>e</u>xed <u>c</u>ell-free <u>to</u>tal <u>R</u>NA <u>seq</u>uencing). DETECTOR-seq utilized a meticulously tailored set of customized guide RNAs to remove large amounts of unwanted RNAs (i.e., fragmented ribosomal and mitochondrial RNAs) in human plasma. Early barcoding strategy was implemented to reduce costs and minimize plasma requirements. Using DETECTOR-seq, we conducted a comprehensive analysis of cell-free transcriptomes in both whole human plasma and EVs. Our analysis revealed discernible distributions of RNA types in plasma and EVs. Plasma exhibited pronounced enrichment in structured circular RNAs, tRNAs, Y RNAs, and viral RNAs, while EVs showed enrichment in mRNAs and srpRNAs. Functional pathway analysis highlighted RNA splicing-related ribonucleoproteins (RNPs) and antimicrobial humoral response genes in plasma, while EVs demonstrated enrichment in transcriptional activity, cell migration, and antigen receptor-mediated immune signals. Our study indicates the comparable potential of cfRNAs from whole plasma and EVs in distinguishing cancer patients (i.e., colorectal and lung cancer) from healthy donors. And microbial cfRNAs in plasma showed potential in classifying specific cancer types. Our comprehensive analysis of total and EV cfRNAs in paired plasma samples provides valuable insights for determining the need for EV purification in cfRNA-based studies. We envision the cost-effectiveness and efficiency of DETECTOR-seq will empower transcriptome-wide investigations in the fields of extracellular vesicles and liquid biopsy.

## Introduction

In recent years, liquid biopsy has emerged as a non-invasive approach for assessing circulating biomarkers in various body fluids, enabling the monitoring of physiologic and disease states [1]. Cell-free RNAs (cfRNAs), given their virtue of being highly dynamic, hold great potential to reflect the pathophysiological processes, thus offering unique opportunities for disease monitoring. Previous reports have suggested that cfRNAs are packaged into various extracellular complexes, such as extracellular vesicles (EVs, including micro-vesicles and exosomes) and non-vesicular ribonucleoproteins (RNPs) [2]. Due to the protection of EV, RNA binding proteins, and/or their self-structures, cfRNAs are capable of being stably present in human bloodstream [3]. While previous studies have predominantly focused on total cfRNAs [4–6] or EV [7–9] cfRNAs in plasma, the transcriptional differences between these two entities remain poorly understood.

Efforts to characterize cfRNAs initially centered around small RNAs like microRNAs (miRNAs) because of the nature of RNA degradation and fragmentation in biofluids. However, miRNAs represent only a small proportion of the human transcriptome [10]. Consequently, investigations have expanded to encompass a broader range of cfRNA species, including messenger RNAs (mRNAs), long non-coding RNAs (lncRNAs), and circular RNAs (circRNAs) [4–7, 11]. These cell-free long RNA species (>50 nt) have relatively low concentrations in human blood due to the presence of RNases, and they are typically fragmented (∼50–200 nucleotides) with incomplete RNA ends [12]. Conventional small RNA-seq approaches, which rely on ligating sequencing adapters based on the 5’ phosphate (5’ P) and 3’ hydroxyl (3’ OH) ends of RNA, are inadequate for analyzing these fragmented cfRNAs [13].

Recently, several sequencing approaches have been developed to profile cell-free long RNA fragments in plasma or EVs. Phospho-RNA-seq integrates T4 polynucleotide kinase into ligation-based TruSeq small RNA-seq, enabling the recovery of mRNA and lncRNA fragments lacking 5’ P and/or 3’ OH ends. However, the libraries generated by Phospho-RNA-seq contain high proportions of ribosomal RNAs (rRNAs) and Y RNAs, limiting the capacity to detect other informative RNA species [12]. Another method, SILVER-seq, captures both small and long cfRNAs from extremely low-input serum samples [14]. However, substantial DNA contamination seemed to be an issue of SILVER-seq [15]. Recently, SMARTer stranded total RNA-seq (hereafter called SMARTer-seq) has been employed in several cfRNA studies [4–7], utilizing a proprietary ZapR and R-probes to deplete unwanted ribosomal sequences [16, 17]. However, as a commercial kit, SMARTer-seq is not specifically optimized for cfRNA library preparation from plasma and is cost-inefficient. Overall, the current cfRNA sequencing approaches were hindered by unwanted RNAs, DNA contamination, and high cost.

Current targeted depletion strategies for unwanted RNAs, such as RiboMinus kits (Thermo Fisher Scientific), Ribo-Zero technology (Illumina), and RNase H-mediated digestion of RNA:DNA hybrids [16], primarily operate at the RNA level and require relatively intact RNA molecules. Consequently, these methods are unsuitable for low-input and fragmented cfRNA samples. In contrast, the Cas9-mediated targeted DNA cleavage technique, also known as DASH (depletion of abundant sequences by hybridization) [18], provides the capability to selectively cleave cDNA molecules derived from rRNAs during the double-stranded DNA stage after library amplification. Notably, this method only requires the design of a set of specific single-stranded guide RNAs (sgRNAs) to direct Cas9 cleavage of undesirable sequences. Therefore, CRISPR-Cas9 presents a highly advantageous approach for the targeted removal of over-represented sequences in the libraries of low-input and fragmented cfRNA samples derived from plasma and EVs.

In this study, we present an optimized cfRNA sequencing method, DETECTOR-seq (<u>de</u>pletion-assisted multipl<u>e</u>xed <u>c</u>ell-free <u>to</u>tal <u>R</u>NA <u>seq</u>uencing), which utilizes early barcoding and CRISPR-Cas9 to reduce costs and deplete highly abundant, fragmented rRNAs and mitochondrial RNAs (mtRNAs) in human plasma. Subsequently, we used DETECTOR-seq to investigate 113 plasma cfRNA samples (including 61 plasma total RNA and 52 EV RNA libraries), derived from healthy donors, lung and colorectal cancer patients. To the best of our knowledge, this study is the first to compare paired total and EV-selected transcriptomes in the same plasma samples, suggesting their distinct signatures and different utilities in cancer liquid biopsy.

## Results

### Development of DETECTOR-seq to profile cell-free transcriptome

The sequencing of cfRNAs in plasma and extracellular vesicles (EVs) usually meets the following obstacles. First, consistent with previous reports [10], we observed that plasma cfRNAs were degraded with a fragment length of <200 nucleotides **(Figure 1A)**. These fragmented cfRNAs are hard to be detected by RNA-seq protocols based on ligation techniques requiring intact RNA ends. Second, ribosomal RNAs (rRNAs) and mitochondrial RNAs (mtRNAs) accounted for ∼92% of all clean reads (reads after removing adapters and filtering low-quality reads), while messenger RNAs (mRNAs) and long non-coding RNAs (lncRNAs) collectively made up only a small fraction (∼4%) of cell-free transcriptome **(Figure 1B)**. It is worth noting that microbe-derived RNAs can also be detected in human plasma with a relatively small fraction (∼0.4%) **(Figure 1B)**. The high fractions of rRNAs and mtRNAs hamper the detection of other informative RNA species. And they are fragmented into pieces in plasma, making them hard to be removed **(Figures 1C, D)**. Third, cfRNAs are usually in the range of hundred picograms to several nanograms per ml of human plasma [14], which can be easily lost and contaminated during purification and amplification. Furthermore, low cfRNA input usually requires 20-24 PCR amplification cycles for library preparation, which produces a high duplication ratio of raw reads. Meanwhile, DNA contamination ignorable in conventional RNA-seq is often over-amplified, causing a big issue in cfRNA-seq [15].

**Figure 1.**
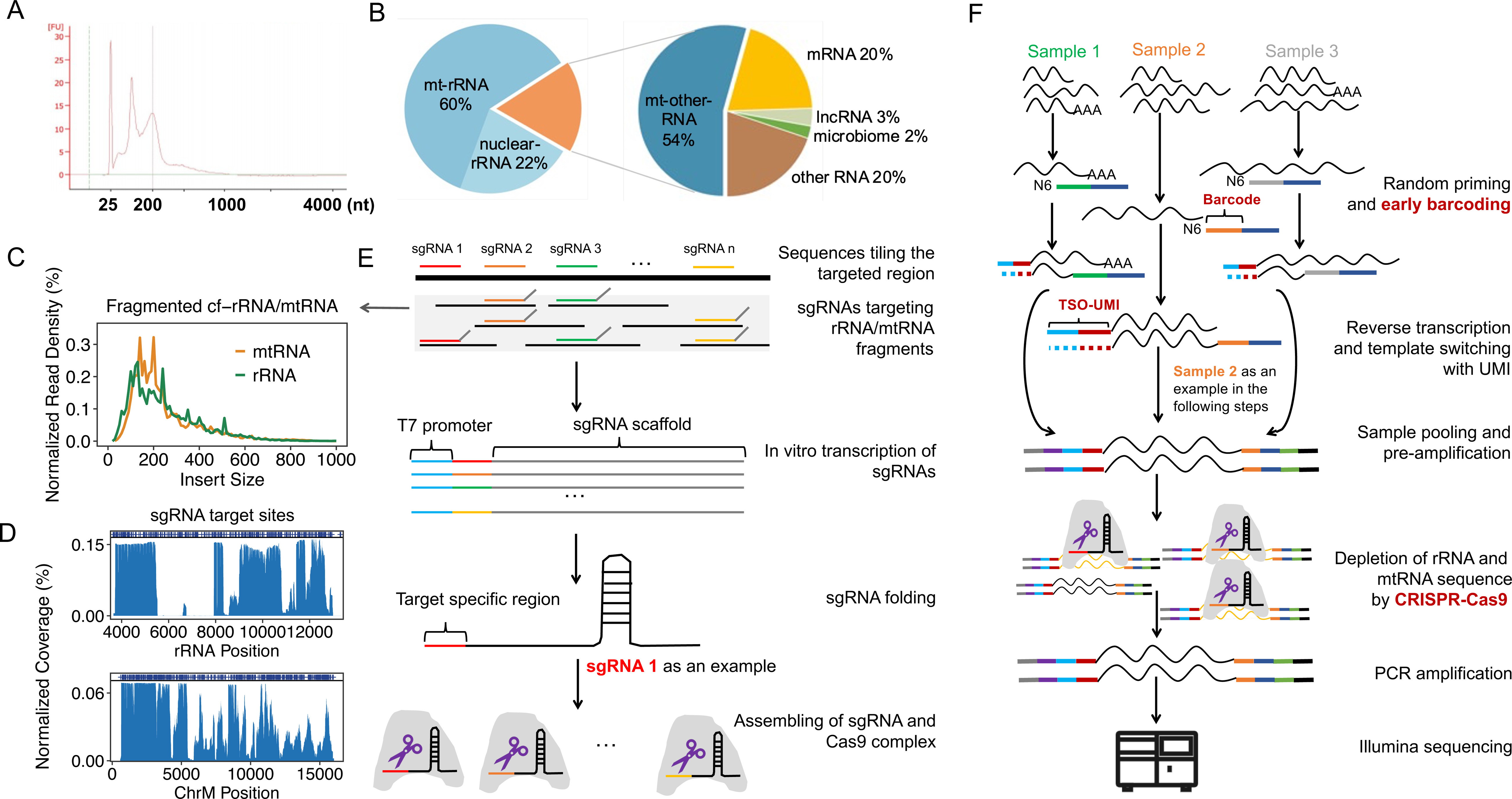
Depletion-assisted multiplexed cell-free total RNA sequencing. **(A)** Bioanalyzer trace of cfRNA fragment lengths in a human plasma sample. (**B)** The relative proportion of reads for various RNA biotypes detected by total RNA sequencing averaged by three human plasma samples. **(C)** Distribution of reads’ insert size for the fragmented rRNAs and mtRNAs, derived from the above sequencing data. **(D)** Distribution of reads’ coverage. Blue bars on top represent sgRNA target sites. **(E)** The designed sgRNAs tiling the fragmented rRNA and mtRNA sequences. **(F)** Schematic overview of DETECTOR-seq workflow. First, cfRNAs are reverse transcribed with random primers and TSO. Sample barcodes and UMIs are introduced during this step. Second, after calibrating input amounts, samples are pooled and pre-amplified. Third, cDNAs of rRNAs and mtRNAs are depleted by CRISPR-Cas9. Subsequently, DETECTOR-seq library is further amplified, then sequenced on an Illumina platform. rRNA: ribosomal RNA; mtRNA: mitochondrial RNA; TSO: template switching oligo; UMI: unique molecular identifier.

To improve the efficiency and reliability of cfRNA detection, we developed DETECTOR-seq (<u>de</u>pletion-assisted multipl<u>e</u>xed <u>c</u>ell-free <u>to</u>tal <u>R</u>NA <u>seq</u>uencing) to profile cell-free transcriptome in human plasma **(Figures 1E, F)**. DETECTOR-seq captures fragmented cfRNAs with unbiased random priming and template-switching. Then, we adapted and modified a previously described method termed CRISPR/Cas9–based Depletion of Abundant Species by Hybridization (DASH) [18] to remove the abundant sequences derived from ribosomal and mitochondrial RNAs in the complementary DNA (cDNA) library. In this step, guide RNAs (sgRNAs) in the CRISPR-Cas9 are specifically optimized for human plasma cfRNAs **(Supplementary Figures 1,2)**, covering the fragmented rRNA and mtRNA sequences **(Figures 1D, E)**. The sgRNAs are in vitro transcribed using T7 RNA polymerase, then bind with Cas9 nuclease to form ribonucleoprotein (RNP) complex and induce site-specific cleavage with the endonuclease activity of Cas9 **(Figure 1E)**, thus preventing further amplification of cDNAs derived from rRNAs and mtRNAs in the final sequencing library. Meanwhile, DETECTOR-seq utilizes early barcoding during reverse transcription. The multiplexed library will cope with low content of plasma cfRNAs, and reduce experimental time and cost as well. It is also worth mentioning that unique molecular identifiers (UMIs) are added to every sequence in the reverse transcription step, hence DETECTOR-seq is capable of removing PCR duplicates to avoid RNA quantification bias **(Figure 1F)**. In addition, we also optimized cfRNA extraction **(Supplementary Figure 3)** and residual DNA digestion **(Supplementary Figure 4)** protocols.

### Analytical validation demonstrating superior performance of DETECTOR-seq

To examine whether DETECTOR-seq can deplete the unwanted rRNA and mtRNA sequences effectively and specifically, we split a single plasma sample into two equal aliquots for experimental conditions of untreated versus depleted, with six biological replicates. In the untreated samples, reads mapped to rRNAs and mtRNAs collectively represented ∼94% of all mapped reads. After CRISPR-Cas9 treatment, these unwanted sequences were decreased to only ∼15% of mapped reads, only about one-sixth of the untreated ones **(Figure 2A)**. By comparing untreated and depleted aliquots, we observed evident decreases in the normalized coverage of rRNAs and mtRNAs **(Figure 2B)**. Meanwhile, the expression levels of detected genes other than rRNAs and mtRNAs between the untreated and depleted aliquots were well correlated, indicating minimal off-target effect (Pearson correlation, R = 0.92, *P*-value < 2.2×^-16^; **Figure 2C**). By comparing the cfRNA expression profiles obtained from DETECTOR-seq and SMARTer-seq, we found that the expression levels of detected genes using these two methods were also well correlated (Pearson correlation, R = 0.90, *P*-value < 2.2×^-16^; **Figure 2D**). In summary, the above results demonstrate the efficient and specific depletion of unwanted sequences in DETECTOR-seq.

**Figure 2.**
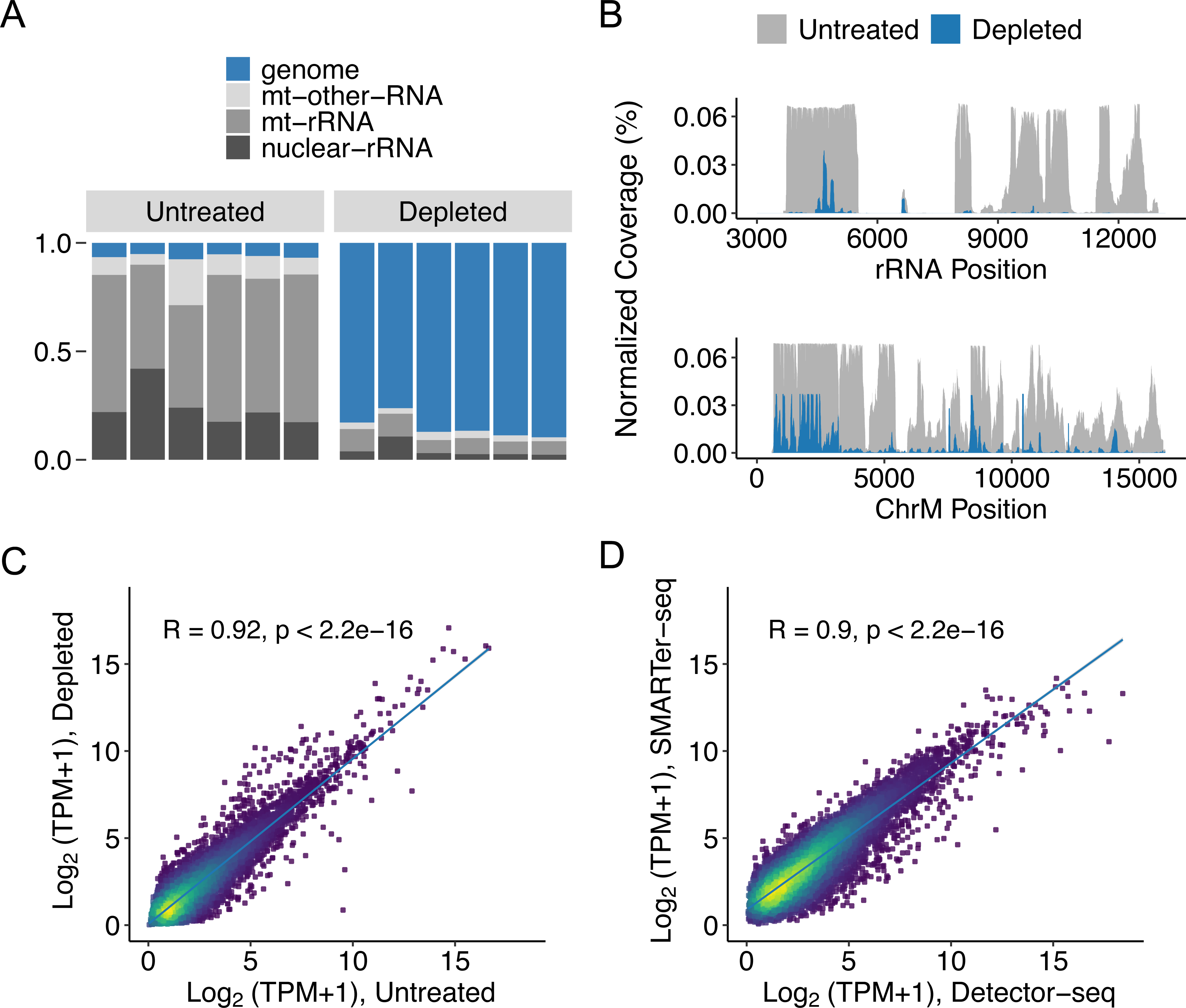
Efficient and specific depletion of rRNA and mtRNA sequences. **(A)** The read distributions and **(B)** coverages of untreated and rRNA/mtRNA-depleted DETECTOR-seq libraries. Read coverage was normalized to total mapped reads. Pearson correlation of cfRNA expression levels between **(C)** untreated and rRNA/mtRNA-depleted DETECTOR-seq libraries, and **(D)** DETECTOR-seq versus SMARTer-seq. TPM: transcripts per million mapped reads (rRNA/mtRNA reads were removed).

To further evaluate the performance of DETECTOR-seq, we prepared cfRNA libraries in a 3-plex, 4-plex, or 5-plex manner determined by RNA concentrations. The total read numbers of different barcoded samples in one multiplexed pool were relatively uniform, varying less than 1.5-fold in the 3- and 4-plex samples and less than 3-fold in the 5-plex samples **(Supplementary Figure 5A)**. In addition, the UMI strategy in DETECTOR-seq retained significantly more reads than the non-UMI approach after duplicated reads were removed **(Supplementary Figure 5B)**. And a sharp edge of reads’ distribution across exon-intron splice junctions suggested that the majority of DNA contamination was effectively removed **(Supplementary Figure 5C)**. To evaluate the impact of plasma input volume on the number of detected genes, we sequenced cfRNAs with 200, 400, 600, 800, and 1000 μL of plasma aliquots from the same individual with five biological replicates. Around 4000 genes were detected with the minimum (i.e., 200 μL) volume. The detected gene number linearly increased until a plateau between 800 and 1000 μL, suggesting the detected genes would be saturated after 1 mL of plasma **(Supplementary Figure 5D)**. While highly correlated cfRNA expression levels were observed within technical triplicates (R1-R3), the correlations were slightly decreased between biological triplicates (N1-N3) **(Supplementary Figure 5E)**. Furthermore, based on ERCC RNA Spike-In Mix, we found a high correlation between expected and observed levels of transcript abundance (Pearson correlation, R = 0.91, *P*-value < 2.2×^-16^; **Supplementary Figure 5F**). These results not only demonstrate DETECTOR-seq’s high accuracy and reproducibility but also suggest its capability of capturing subtle differences in cfRNA profiles between different individuals.

Then, we randomly subsampled a dataset (n=24) of DETECTOR-seq for saturation analyses of detected UMIs (transcripts) and genes. Although the detected UMIs kept increasing when more reads in 1ml plasma were sequenced **(Supplementary Figure 5G)**, the detected gene numbers were quickly saturated at approximately 5 million genome-aligned reads **(Supplementary Figure 5H)**. These results indicate that DETECTOR-seq achieves saturation of cfRNA detection at a low sequencing depth.

### Better contamination control and cost-effectiveness of DETECTOR-seq than other cfRNA-seq methods

We benchmarked the performance of DETECTOR-seq compared to three other cfRNA-seq methods, including Phospho-RNA-seq [12], SILVER-seq [14], and SMARTer-seq [19]. The number of samples used in the comparison was listed in **Supplementary Table 5**. Within the total genome-aligned reads, DETECTOR-seq and SMARTer-seq had comparable ratios of exonic reads (∼70%), while those of SILVER-seq and Phospho-RNA-seq were under 40% **(Figure 3A)**. The lower ratio of exonic reads for SILVER-seq was presumably due to severe DNA contamination according to a previous report [15]. We also visualized the read coverage across exon boundary sites flanked upstream and downstream by 50 bp, where DETECTOR-seq and SMARTer-seq showed more evident decreases of read coverage from exon to intron/intergenic region than SILVER-seq and Phospho-RNA-seq **(Figure 3B)**. As far as we know, all of the four cell-free RNA-seq methods should preserve the strand specificity of RNAs. Thus, the enrichment of exons’ sense over antisense reads of DETECTOR-seq and SMARTer-seq further confirmed their reads’ quality **(Figure 3C)**. The above results demonstrate that DETECTOR-seq and SMARTer-seq have better DNA contamination control than SILVER-seq. It was worth noting that Phospho-RNA-seq was developed from a small RNA-seq method, and the read coverage across exon boundary sites and the enrichment of exons’ sense over antisense reads may be affected by the read distribution of small RNAs.

**Figure 3.**
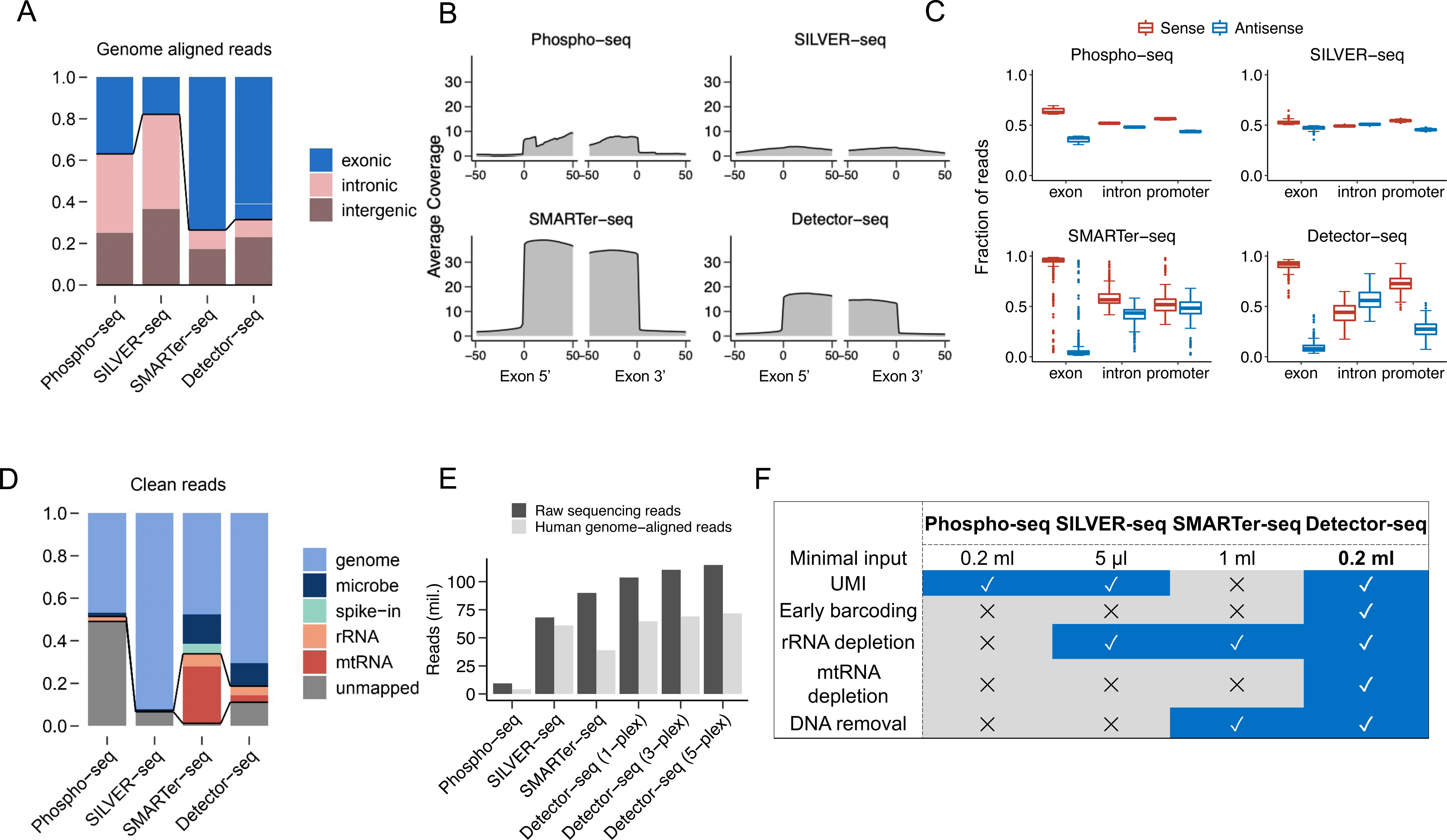
Comparing DETECTOR-seq with other cfRNA-seq methods. **(A)** Average percentages of genome-aligned reads mapping to exonic, intronic, and intergenic regions for four different cfRNA-seq methods. **(B)** Average coverage across all mRNAs’ 5’ and 3’ exon boundary sites flanking upstream and downstream by 50 bp. **(C)** Average percentages of reads located in the sense and antisense strands of mRNAs’ exons, introns, and promoters. **(D)** Average percentages of clean reads (after trimming low-quality and adapter sequences) assigned to different sources. **(E)** Numbers of raw sequencing reads and human genome-aligned reads with a fixed budget of $300 for each method. **(F)** Summary of key techniques used in the four cfRNA-seq approaches. Numbers of used samples: Phospho-seq: 15; SILVER-seq: 128; SMARTer-seq: 373; DETECTOR-seq: 113.

In addition, we showed that DETECTOR-seq displayed a higher ratio of reads mapped to human genome (∼71%) than those of SMARTer-seq (∼48%) because DETECTOR-seq removed mitochondrial RNAs more efficiently than SMARTer-seq **(Figure 3D)**. Furthermore, because of its early barcoding and multiplexing strategy, DETECTOR-seq can produce more raw reads and genome-aligned reads than the other cfRNA-seq approaches **(Figure 3E, Supplementary Figure 6)**. Cost details were explained in Supplementary Tables 6 and 7. Overall, by summarizing and comparing key characteristics of these approaches **(Figure 3F)**, we collectively demonstrate that DETECTOR-seq has better contamination control and more efficient cost than the other cfRNA-seq methods.

### Distinct human and microbial RNA signatures in plasma versus extracellular vesicles

Subsequently, we employed DETECTOR-seq to conduct pairwise investigations of total cfRNAs and EV cfRNAs in human plasma **(Figure 4A)**. A proportion of cfRNAs are enclosed inside EVs such as MVs and exosomes [20]. Meanwhile, it is also reported that a significant proportion of cfRNAs are not within EVs but associated with proteins to form non-vesicular RNPs [21]. Although both plasma total cfRNAs [4–6] and EV cfRNAs [7, 9] have been used in liquid biopsy studies, a pairwise comparison of their distinct signatures and utilities has not been conducted yet.

**Figure 4.**
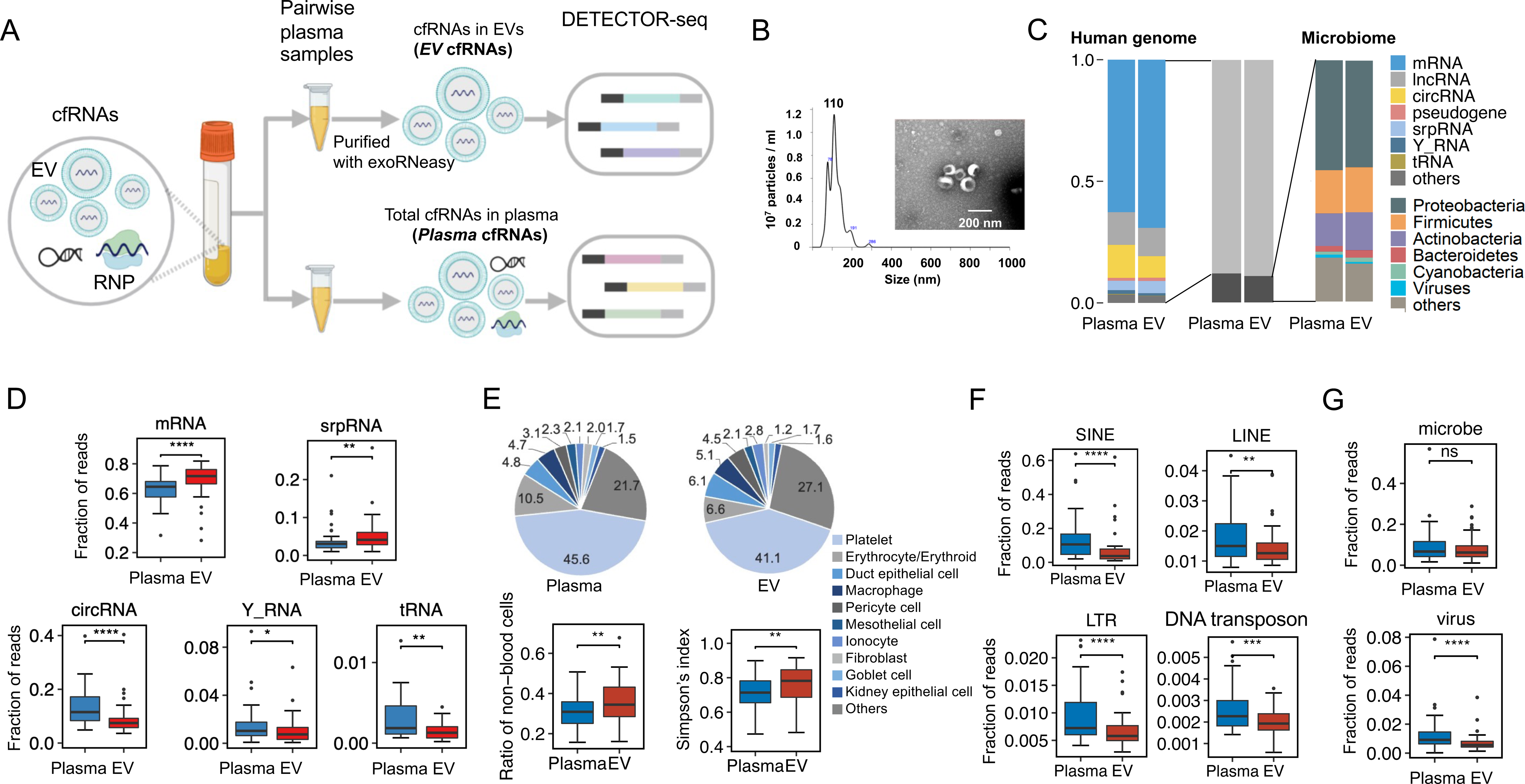
Distinct human and microbial RNA signatures in *Plasma* versus *EV*. **(A)** Illustration of sequencing *Plasma* cfRNAs and *EV* cfRNAs in paired plasma samples. (B) Plasma extracellular vesicles were characterized by nanoparticle tracking analysis and transmission electron microscopy (scale bar represents 200 nm). **(C)** Distribution of reads mapped to human genome and microbiome in *Plasma* and *EV* cfRNA datasets. Left: RNA spectrum mapping to human genome; Right: relative abundance of reads aligned to different phyla. **(D)** Differential human RNA species between *Plasma* and *EV* cfRNAs. **(E)** Pie charts show the average fractional contributions of various cell types to the *Plasma* and *EV* transcriptomes. Box plots show the diversity of cell type contributions to the *Plasma* and *EV* transcriptomes measured by the ratio of non-blood cells and Simpson’s index. **(F)** Boxplots represent the enrichment of cfRNAs derived from transposable elements in *Plasma* cfRNAs compared to *EV* cfRNAs. The different transposable element categories, including short interspersed elements (SINEs), long interspersed elements (LINEs), long interspersed elements with long terminal repeats (LTR), and DNA transposons, are represented. **(G)** The fractions of reads aligned to microbe and virus. *Plasma*: 44 samples; *EV*: 44 samples (all samples paired). ****: *P*-value < 0.0001, **: *P*-value < 0.01, *: *P*-value < 0.05, Wilcoxon rank sum test, two-tailed.

In this study, a total of 139 plasma cfRNA samples were sequenced, which included samples obtained from healthy donors as well as patients with lung cancer and colorectal cancer (**Supplementary Figure 7, Supplementary Table 10**). EVs were purified using a membrane affinity column, concentrating particles predominantly within the size range of 50 to 200 nm, with a peak around 110 nm. Morphological examination using transmission electron microscopy confirmed the presence of the characteristic cup-shaped structure commonly associated with EVs **(Figure 4B)**. After conducting quality control (QC) procedures on the RNA samples and sequencing data, a total of 113 datasets passed the QC criteria (**Supplementary Figures 7-9**). Out of these 113 datasets, 61 were derived from total cfRNA-seq of plasma, while 52 were obtained from EV cfRNA-seq of plasma. Among them, 44 datasets were paired, meaning they originated from the same plasma samples. In the following description, total cfRNA-seq of plasma and EV cfRNA-seq of plasma will be abbreviated to *Plasma* cfRNA and *EV* cfRNA, respectively.

From a general view, there was a high degree of similarity between *Plasma* and *EV* cfRNAs, with ∼90% of aligned reads mapping to human genome and ∼10% mapping to microbe genomes **(Figure 4C)**. For human cfRNAs, mRNA, lncRNA, and circRNA were the major RNA types. For microbial cfRNAs, the most abundant phylum was *Proteobacteria*, followed by *Firmicutes* and *Actinobacteria*. The human and microbial RNA compositions resembled previous reports [19, 22].

In addition, distinctive signatures were revealed for the first time by our pairwise comparison between *Plasma* (n = 44) and *EV* (n = 44) cfRNAs (all samples were paired). We first observed that *Plasma* cfRNAs had more short fragments (20∼100 nt), while *EV* cfRNAs had more long fragments (>100 nt) **(Supplementary Figure 10)**. We also observed that structured tRNAs, Y RNAs, and circRNAs were enriched in *Plasma* cfRNAs, while mRNAs and signal recognition particle RNAs (srpRNAs) were enriched in *EV* cfRNAs **(Figure 4D)**. These findings align with a previous study that reported a significant enrichment of tRNA and Y RNA fragments in extracellular RNPs [2]. Moreover, we also found that the relative abundance of circRNAs was slightly higher in *Plasma* cfRNAs than *EV* cfRNAs (median 13.6% vs. 8.8%, *P*-value < 0.0001, Wilcoxon rank sum test; **Figure 4D**, **Supplementary Figure 11)**, perhaps due to its circle-like structure resisting degradation outside of EVs. We totally identified 13 circRNAs differentially enriched in *Plasma* versus *EV* cfRNAs. Only one of them, hsa_circ_0048555, was enriched in EVs **(Supplementary Figure 12)**. Reads mapped to the back-spliced junction were used to calculate the enrichment.

A recent study provided a framework to infer cell types of origin of the cell-free transcriptome [23]. We utilized this method and found a high similarity of the cell types of origin between *Plasma* and *EV* transcriptomes **(Figure 4E)**. Platelets and erythrocytes were inferred as the major origins for both *Plasma* and *EV* cfRNAs, which was in agreement with the previous study [23]. Intriguingly, we found non-blood cells contributed more to *EV* cfRNAs than to *Plasma* cfRNAs (*P*-value < 0.01, Wilcoxon rank sum test; **Figure 4E**). Therefore, the diversities of cell types of origin (measured by Simpson’s index) of *EV* cfRNAs were slightly higher than those of *Plasma* cfRNAs (0.75 vs. 0.70, *P*-value < 0.01, Wilcoxon rank sum test; **Figure 4E**).

A noteworthy discovery has been made regarding the presence of RNAs originating from transposable elements and other repetitive elements in the cell-free transcriptome [24]. In our current investigation, we provide evidence demonstrating a significant enrichment of cfRNAs derived from transposable elements (TEs) in *Plasma* cfRNAs compared to *EV* cfRNAs. These transposable elements include short interspersed elements (SINEs), long interspersed elements (LINEs), long interspersed elements with long terminal repeats (LTR), and DNA transposons **(Figure 4F)**.

We also identified distinct microbe genera in *Plasma* and *EV* cfRNAs **(Supplementary Figure 13)**. While there was no significant difference in the ratio of microbial reads between *Plasma* and *EV* cfRNAs, we did observe a significant increase in cfRNAs mapped to viral genomes in *Plasma* cfRNAs **(Figure 4G)**. Meanwhile, viruses such as *Senecavirus*, *Cheravirus*, *Orthopoxvirus*, *Tenuivirus*, and *Rhadinovirus* were enriched in *Plasma* cfRNAs, while *Intestinimonas*, *Mordavella*, and *Jonquetella* were enriched in *EV* cfRNAs **(Supplementary Figure 13)**. In summary, the above comparison results have revealed distinct molecular characteristics between *Plasma* and *EV* cfRNAs in terms of fragment size, RNA species, cell types of origin, TE RNAs and microbe genera.

### Functional pathways and sequence motifs of selective *Plasma* and *EV* cfRNAs

To find selective functions and motifs of cfRNAs in EVs, we identified 545 selectively distributed RNAs showing significantly differential abundance between *Plasma* and *EV* transcriptomes (|Fold-change| >1 and FDR < 0.1; **Figure 5A, Supplementary Figure 14)**. Among them, 271 cfRNAs were enriched in *Plasma*, while 274 cfRNAs were enriched in *EVs*. We investigated the functional roles and biological pathways of these selective cfRNAs **(Figure 5B, Supplementary Figure 14)**. Based on KEGG pathway enrichment analysis, we found that the selective RNAs elevated in *Plasma* were significantly enriched in terms associated with RNA splicing, RNP (e.g., mRNA 5’ splice site recognition, U1 snRNP, spliceosomal snRNP complex and Sm-like protein family complex), antimicrobial and innate immune responses. Meanwhile, the selective RNAs that were enriched in *EVs* were primarily associated with DNA binding transcription factor activity, focal adhesion, cell-substrate junction, and T cell receptor signaling immune pathway. Notably, we also found different immune pathways enriched in the selective cfRNAs of *Plasma* versus *EVs* **(Figure 5B, Supplementary Figure 15)**. The organ or tissue-specific immune response and antimicrobial humoral response immune signaling pathways are enriched in *Plasma* cfRNAs, while defense response to other organisms, Fc receptor signaling pathway, T cell receptor signaling pathway are enriched in *EV* cfRNAs **(Supplementary Figure 15)**.

**Figure 5.**
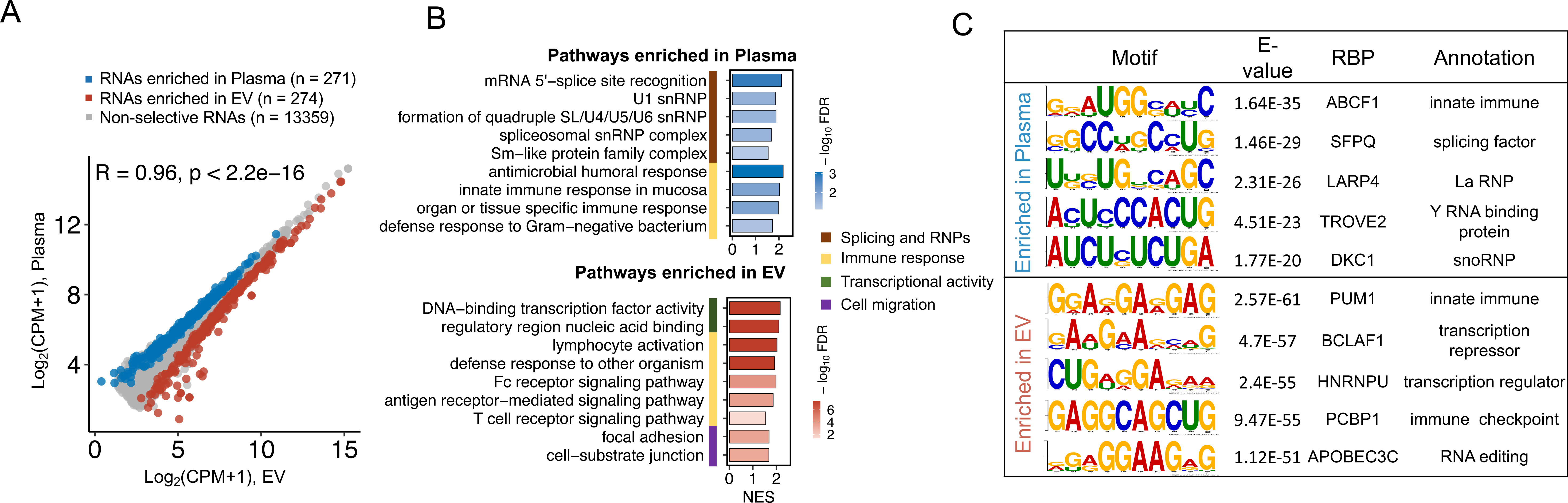
Distinct functional pathways, motifs, and binding proteins of the selective *Plasma* and *EV* cfRNAs. **(A)** Definition of the selective cfRNAs enriched in *Plasma* or *EV*. Cutoff: |Fold-change|>1 and FDR<0.1. **(B)** Top enriched GO pathways of the selective cfRNAs. **(C)** Top enriched motifs and their corresponding RNA binding proteins (RBPs) of the selective cfRNAs. *Plasma*: 44 samples; *EV*: 44 samples (all samples paired).

We further investigated sequence motifs and their associated RNA binding proteins (RBPs) for the selective cfRNAs **(Figure 5C, Supplementary Figure 16)**. And we found that the selective cfRNAs enriched in *Plasma* contained binding motifs/sites for ABCF1, a protein that plays a role in innate immune response [25]; SFPQ, a splicing factor; LARP4, a La RNP; TROVE2, a Y RNA binding protein; and DKC1, a snoRNP. Meanwhile, the selective cfRNAs enriched in *EVs* contained binding motifs/sites for PUM1, a protein that participates in human innate immune response [26]; BCLAF1, a transcription factor; HNRNPU, a transcription suppressor; PCBP1, a previously reported immune checkpoint [27]; APOBEC3C, an RNA editing enzyme. These enriched motifs and their associated RBPs were consistent with the biological functions of the selective cfRNAs revealed above.

### Specific cancer-related signals in *Plasma* and *EV* cfRNAs

Next, we compared the potential of *Plasma* cfRNAs and *EV* cfRNAs to discriminate between cancer patients and healthy individuals in a proof-of-concept cohort. We sequenced cfRNAs in the plasma samples of lung cancer (LC, Plasma n = 19, EV n = 19, 18 of them paired) and colorectal cancer (CRC, Plasma n = 23, EV n = 19, 19 of them paired) patients **(Supplementary Figure 7)**. To maximize the sample size, we merged CRC and LC together as a combined cancer group. Based on differential expression analysis between this combined cancer group (Plasma n = 42, EV n = 38, 37 of them paired) and normal controls (NC, Plasma n = 19, EV n = 14, 7 of them paired) using the criteria of |log2fold-change|>1 and FDR<0.05, we defined a set of cancer-relevant cfRNAs in both *Plasma* and *EVs* **(Supplementary Figure 17)**. Interestingly, when we intersected the cancer-relevant cfRNAs and selective cfRNAs mentioned above, we found that cancer-relevant cfRNAs accounted for 59.8% (162/271) of the selectively enriched cfRNAs in *Plasma*, whereas they only represented 6.9% (19/274) of the selectively enriched cfRNAs in *EVs*. Therefore, cancer-relevant cfRNAs appear to be more enriched in *Plasma*’s selective cfRNA fraction. **(Figure 6A).** We also found that enriched functions of these cancer-relevant *Plasma* cfRNAs were termed as RNA splicing, snRNP signals, etc **(Figure 6B)**, which were consistent with the enriched pathways of *Plasma* cfRNAs revealed in **Figure 5B**.

**Figure 6.**
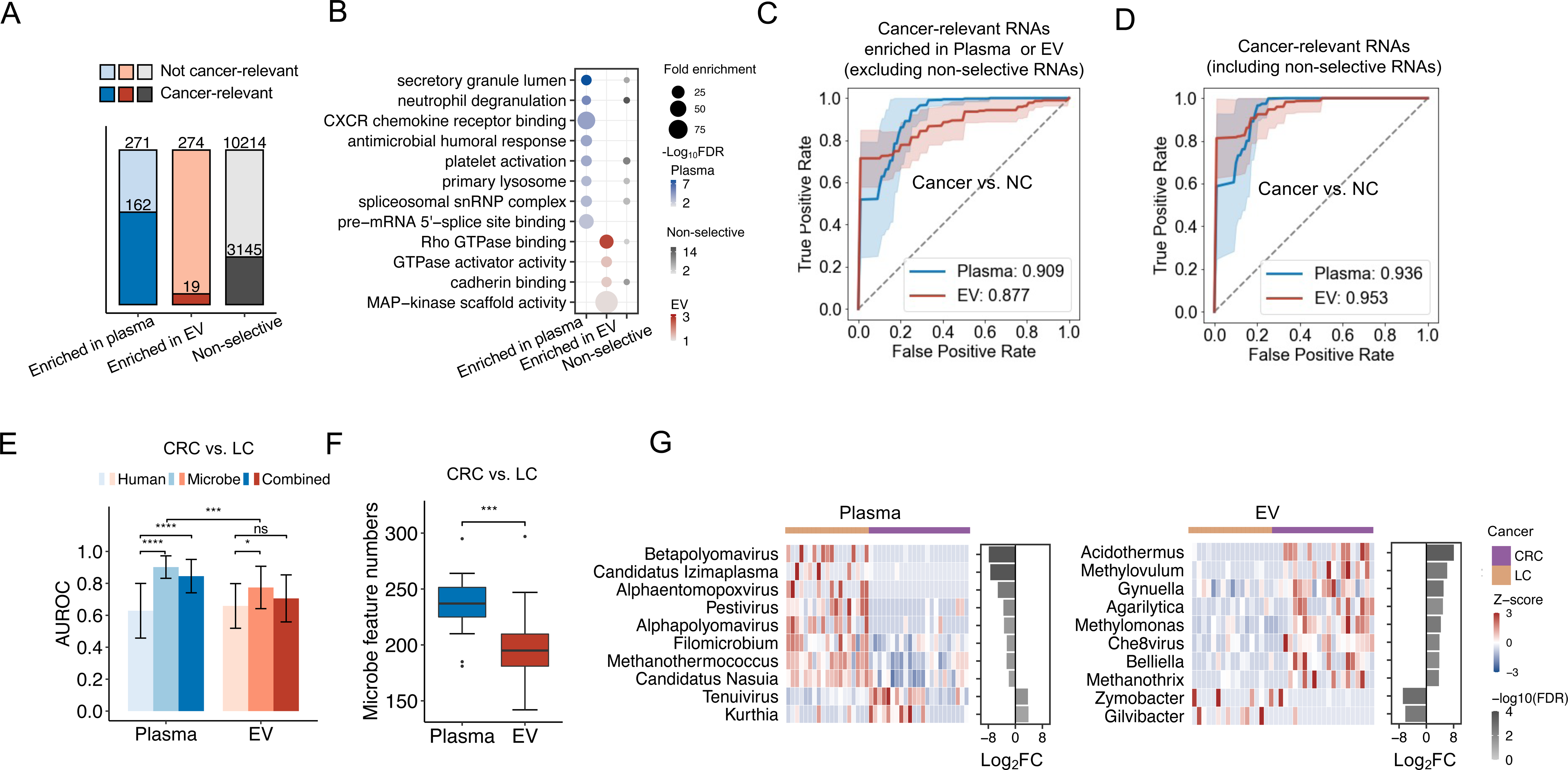
Cancer-relevant cfRNA signatures in *Plasma* and *EV*. **(A)** Cancer-relevant ones (differentially expressed between cancer patients and normal controls, |log2fold-change|>1 and FDR<0.05) in the selective and non-selective human cfRNAs. Cancer: colorectal cancer (CRC) and lung cancer (LC); NC: normal control. **(B)** Enriched GO terms related to cancer-relevant human cfRNAs. Performances (average of 20 bootstrap procedures) of cancer-relevant human cfRNAs distinguishing cancer patients from normal controls when excluding **(C)** and including **(D)** non-selective cfRNAs. **(E)** AUROCs of cancer type classification (CRC vs. LC) using human- or microbe-derived reads in *Plasma* and *EV* cfRNAs. **(F)** Numbers of microbial features (genus) with significantly differential abundance (|log2fold-change|>1 and FDR<0.1) between CRC and LC in 20 bootstrap procedures. **(G)** Distinct cancer type-specific microbial features (genus) identified in *Plasma* and *EV* cfRNAs. Heatmaps show z-scores of the abundance levels of these microbial RNA features; bar plots illustrate their average log2FCs and FDRs between CRC and LC. FC: fold-change; FDR: false discovery rate. ****: *P*-value < 0.0001, ***: *P*-value < 0.001, *: *P*-value < 0.05, Wilcoxon rank sum test, two-tailed. CRC samples: *Plasma* (n=23), *EV* (n=19), 19 of them paired; LC samples: *Plasma* (n=19), *EV* (n=19), 18 of them paired; NC samples: *Plasma* (n=19), *EV* (n=14), 7 of them paired.

Based on these selectively distributed cancer-relevant cfRNAs, we endeavored to discriminate cancer patients from NCs. Although the selective cfRNAs in *Plasma* performed slightly better than those in *EVs* (average AUROC: 0.909 versus 0.877, **Figure 6C, Supplementary Figure 18)**, comparable performances were observed between *Plasma* and *EV* cfRNAs when a large number of non-selective cfRNAs were included as well (average AUROC: 0.936 versus 0.953, **Figure 6D, Supplementary Figure 19**). Collectively, these results imply that the purification of EV can reveal distinct cancer signals, but it has a very subtle effect on the accuracy of detection of cancer patients from healthy controls.

We further assessed the potential of cfRNAs (human-derived only) in *Plasma* and *EV* for classifying CRC from LC. Initially, neither cfRNAs in *Plasma* or *EV* exhibited strong classification potential (average AUROC: 0.628 versus 0.659, **Figure 6E, Supplementary Figure 20)**. A recent study revealed that microbe-derived cfRNAs in human plasma reflect cancer-type-specific information [19]. Based on the RNA abundance of the contamination-filtered microbe genera, we found the microbial cfRNAs improved the classification of cancer types for both *Plasma* and *EV* cfRNAs (average AUC: 0.898 versus 0.772, **Figure 6E, Supplementary Figure 20)**.

Notably, the microbial cfRNAs in *Plasma* performed better than those in *EV*. Consistently, we also found more cancer-type-specific features in *Plasma* cfRNAs than in *EV* cfRNAs **(Figure 6F)**. We identified the microbial features recurrently showing differential abundance between CRC and LC in all of the 20 bootstrap samplings. The abundance of top recurrent microbe genera, along with fold-change and false discovery rates were illustrated **(Figure 6G)**. For instance, we observed a higher relative abundance of *Methanothrix* in CRC compared to LC using *EV* cfRNA-seq data. This is consistent with a previous study reporting that *Methanothrix soehngenii* was enriched in gut microbiome of CRC patients [28]. Meanwhile, many cancer-relevant viral RNAs in *Plasma* classified cancer types, consistent with the observation of more viral RNAs detected in *Plasma* than in *EVs* **(Figure 4F)**. For instance, *Plasma* cfRNA-seq data revealed a higher abundance of *Alpha-polyomavirus* and *Beta-polyomavirus*. Supportively, some polyomaviruses were also reported to be detectable in gastrointestinal tract and respiratory aspirates [29]. These findings suggest that microbe-derived cfRNAs in *Plasma*, at least in this small cohort with limited sample size, present promising but yet poorly investigated signatures for specific cancer types. Further validation in larger cohorts is required to establish the clinical utility and significance of these preliminary findings.

## Conclusion and Discussion

### Conclusions

In summary, this study introduced a depletion-assisted cost-effective cfRNA profiling approach, termed DETECTOR-seq, which overcomes challenges associated with low quantity and low quality of fragmented cfRNAs, over-represented rRNAs and mtRNAs, DNA contamination, and high costs. Using DETECTOR-seq, we recapitulated molecular characteristics of *Plasma* and *EV* cfRNAs and identified their distinct human and microbial signatures, thus illustrating the gain and loss of certain cfRNA signals due to EV purification. Our work provides a practical reference for researchers engaged in plasma and EV cfRNA-based liquid biopsy **(Table 1)**. Moreover, we envision that DETECTOR-seq would be a useful tool to facilitate further studies in the fields of extracellular RNA biology and plasma or EV cfRNA-based liquid biopsy, paving the way for advancements in both fundamental research and translational medicine.

**Table 1.** Practical reference for cfRNA-seq in human plasma.

|  | *Plasma* cfRNA | *EV* cfRNA |
| --- | --- | --- |
| **EV Purification** | **No** | **Yes** |
| Cost of plasma volume, experimental time and reagents<sup>1</sup> | relatively less | relatively more |
| Enriched RNA species | circRNA, tRNA, Y RNA | mRNA, srpRNA |
| Enriched microbes | viruses | intestinimonas, etc. |
| Diversity of cell-types-of-origin | relatively low | relatively high |
| TE RNAs<sup>2</sup> | relatively more | relatively less |
| Cancer detection | good (AUC: 0.94) | good (AUC: 0.95) |
| Cancer type-specific microbes | relatively more | relatively less |
| Cancer type classification | relatively good (AUC: 0.90) | relatively poor (AUC: 0.77) |
| <sup>1</sup> Different cost is due to the EV purification step. |  |  |
| <sup>2</sup> Cell-free RNAs derived from transposable elements. |  |  |

### Technologies utilized and optimized in DETECTOR-seq

Plasma cell-free transcriptome remains challenging to study owing to the low quantity and quality of fragmented cfRNAs [11]. Over-represented rRNA and mtRNA species [12], DNA contamination [15], and high cost are still the major issues of cfRNA sequencing. Multiple technologies were included in DETECTOR-seq to address these issues **(Figure 4F)**. First, DETECTOR-seq captures fragmented cfRNAs with random priming and template-switching strategies, which have been proven to be highly efficient in single-cell RNA-seq [30]. Second, the early barcoding protocol of DETECTOR-seq enables us to prepare cfRNA libraries in a multiplexed manner, thus reducing the volume of required plasma and experimental costs. In fact, DETECTOR-seq is capable of detecting cfRNAs with a low input volume of 0.2 to 1 mL plasma with a 2- to 6-fold cost saving compared to existing approaches. Third, with UMIs tagging to cDNAs of RNA fragments, DETECTOR-seq can accurately quantify the low-quantity cfRNAs. Fourth, by optimizing the procedures of RNA extraction and residual DNA digestion (**Supplementary Figures 3-4**), DETECTOR-seq avoids the potential contamination of genomic DNAs. Fifth, DETECTOR-seq uses CRISPR-Cas9 technology to deplete rRNA and mtRNA sequences. A CRISPR-based depletion strategy, DASH (Depletion of Abundant Sequences by Hybridization) [18] has been utilized in other fields, such as ATAC-seq [31], small RNA-seq [32], bacterial RNA-seq [33], Ribo-seq [34] and single-cell total RNA-seq [35]. Here, we applied this CRISPR-based method to cfRNA sequencing and designed a specific set of sgRNAs for human plasma **(Supplementary Figures 1,2)**. Of note, our sgRNAs target almost the entire length of human rRNAs and mtRNAs, enabling the use of our guides to deplete rRNAs and mtRNAs from any intact or fragmented RNA samples, regardless of the specimen type. This underscores the versatility of our approach beyond plasma samples.

### *Plasma* vs. *EV* in cancer detection and cancer type classification

Researchers have used both *Plasma* cfRNA-seq [4–6] and *EV* cfRNA-seq [7, 20, 36–38] to identify disease biomarkers. By pairwise comparison between *Plasma* and *EV* cfRNA, we found that both of them can distinguish cancer patients from controls with comparable performance. However, cancer types can be better classified with microbe-derived features in *Plasma* cfRNAs than those in *EV* cfRNAs.

### Distinct signatures in *Plasma* vs. *EV* cfRNAs

This study has brought new insights into distinct cfRNA signatures in *Plasma* versus *EVs*. *Plasma* contains miscellaneous cfRNAs released from alive or apoptotic cells, while RNAs in *EV* cargos are considered to be secreted actively by cells for functional roles in intercellular communications [39]. This study revealed distinct biological pathways, enriched motifs, and RBP-binding sites in *Plasma* vs. *EV* cfRNAs. We also found that short RNA fragments (20 to 100 nt) associated with RNPs were enriched in *Plasma* cfRNAs, indicating higher degradation extent of non-vesicular RNAs than those of EV RNAs.

### Limitations of this study

While analyzing paired plasma samples can increase statistical power, it is important to note that the conclusions regarding the comparison of cfRNAs in *Plasma* and *EVs* for cancer differentiation in this study are still preliminary due to the small sample size. These results serve as a proof-of-concept exploration of DETECTOR-seq’s potential for uncovering intriguing insights in real-world clinical samples. Larger-scale cohorts are required to validate these findings and establish their clinical utility. Furthermore, although DETECTOR-seq offers several advantages compared to other approaches, there is room for further improvement. For instance, the efficiency of random priming in DETECTOR-seq is influenced by the fragment length of RNAs, which can introduce bias in the library preparation. And DETECTOR-seq involves several purification steps to eliminate by-products such as empty library constructs, adapter dimers, and excessive primers. These purification procedures tend to retain longer RNA fragments, resulting in the discarding of RNA fragments shorter than 50 nucleotides, along with the by-products. To obtain a complete spectrum of cfRNAs, including both small and long fragments, DETECTOR-seq could be modified by incorporating alternative strategies such as poly(A) tailing [40, 41].

## Materials and Methods

### Cohort design

Seventy-five participants, including patients with colorectal cancer (n=24), lung cancer (n=20), and healthy controls (n=31), were enrolled in this study. Samples were obtained between November 2018 to January 2022. Individuals with colorectal, lung cancer and healthy controls were recruited from Peking University First Hospital. The characteristics of participants in this study were summarized **(Supplementary Table 9)**.

### Sample collection

Peripheral whole blood samples were collected in EDTA-coated vacutainer tubes for each participant. Blood samples of patients with cancer were collected before any treatment of surgery, chemotherapy, or neoadjuvant chemotherapy. Within 2 hours after blood collection, blood samples were centrifuged at 1,900g for 30 min at room temperature. Plasma was separated and then centrifuged at 16,000g for another 10 min at 4 °C to remove cellular debris. All plasma samples were aliquoted and stored at -80 °C until analysis.

### Plasma and extracellular vesicles RNA extraction

Cell-free RNAs (cfRNAs) were extracted from 1 ml of plasma using QIAzol Lysis Reagent (Qiagen, 79306) according to the manufacturer’s instructions. The upper, aqueous phase containing cfRNAs was mixed with 1 volume of ethanol (95-100%) and then added to the Zymo-Spin column (Zymo, R1016) for RNA binding. Samples were subsequently washed, eluted, and treated with DNase I (TaKaRa, 2270A) for 20 min at 37 °C. Following residual DNA digestion, cfRNAs were then purified and concentrated into 6 µl using an RNA Clean and Concentrator-5 kit (Zymo, R1016). Plasma extracellular vesicles (EVs) were purified by a membrane-affinity approach using an exoRNeasy Midi Kit (Qiagen, 77144) following the manufacturer’s instructions [42]. EVs were eluted with 400 µl of elution buffer and characterized by transmission electron microscopy (TEM) and nanoparticle tracking analysis (NTA). For RNA isolation, EVs were lysed on the exoRNeasy column using QIAzol Lysis Reagent, and EV RNAs were extracted and purified using Zymo-Spin column as mentioned above.

### Optimization of cell-free RNA extraction and residual DNA digestion

RNA extraction is one of the most critical steps for low-input RNA-seq. To this end, we compared three cfRNA extraction approaches, including QPCB (<u>Q</u>IAzol lysis, <u>p</u>henol-chloroform extraction, and <u>c</u>olumn <u>b</u>inding), Norgen (Plasma/Serum Circulating and Exosomal RNA Purification Kit), and QPIP (<u>Q</u>IAzol lysis, <u>p</u>henol-chloroform extraction, and isopropanol <u>p</u>recipitation). QPCB was considered the best approach for cfRNA extraction. (Supplementary Figure 3).

In previous reports, DNA contamination has been emphasized as a hinder to the cfRNA study [15]. Therefore, we examined two major residual DNA digestion approaches: On-column vs. In-buffer (On-column: residual cell-free DNA was digested on the spin-column during RNA extraction; In-buffer: DNA was digested in the aqueous buffer after RNA extraction). We observed a significantly higher human genome mapping ratio and exonic read ratio with In-buffer DNA digestion than On-column approach (*P*-value < 0.0001, Wilcoxon rank sum test; **Supplementary Figure 4**), suggesting In-buffer DNA digestion was more effective to a certain extent. DETECTOR-seq was carried out in the following assays with RNAs extracted using QPCB and residual DNA digested with an In-buffer approach unless specified.

### Reverse transcription

Cell-free RNAs were captured using random primers with a unique sample barcode and then reverse transcribed with SMARTScribe reverse transcriptase (Clontech, 639538) and template-switching oligos tagging 8-nt UMI sequences. Sample barcodes were designed in R using the DNABarcodes package [43]. We generated barcodes with a length of 4 nucleotides and a minimum Hamming distance of 3 and filtered self-complementary sequences, triplets, and sequences that have an unbalanced ratio of bases G or C versus A or T. PEG 8000 (Beyotime, R0056-2ml) was used as molecular crowding reagent to further improve the efficiency of reverse transcription reaction [44]. The 20-µl reaction mixture was incubated at 42 °C for 90 min with a heat inactivation step at 70 °C for 10 min. Primers for the reverse transcription of DETECTOR-seq were shown in **Supplementary Table 3**.

### Quantitative PCR analysis

The total abundance level of *Plasma* or *EV* cfRNAs was assessed by amplifying a fragment from the human gene of *ACTB* spanning the exon-exon junction (ACTB-ee). The level of residual DNA contamination was measured by amplifying a short fragment from *ACTB* within intron regions (ACTB-i). We measured the microbiome contamination by the threshold cycle (Ct) value difference of the ACTB-ee and the bacterial 16S ribosomal RNA V4 fragment (16S-V4). The 2.5 μl of cDNA template was amplified in a final volume of 20 μl using the FastFire qPCR PreMix (SYBR Green) (TIANGEN, FP207). Samples with low RNA content (Ct of ACTB-ee > 32) or high DNA contamination (Ct of ACTB-i < 35), or high bacterial contamination (ΔCt (ACTB-ee – 16S-V4) > 5) were excluded for further analysis. We summarized the quality control primers for *Plasma* and *EV* cfRNA samples **(Supplementary Table 1).**

### Design of guide RNAs (gRNAs)

To remove highly abundant sequences (rRNAs and mtRNAs) in the cfRNA library of human plasma, we designed 302 and 315 high-quality single guide RNAs (sgRNAs) specifically targeting the ribosomal and mitochondrial RNA sequences **(Supplementary Figure 1)**. The sgRNAs were selected and filtered by DASHit [45] and Benchling (https://www.benchling.com/crispr) based on the following criteria: 1) off-target score (specificity score) and on-target score (sensitivity score); 2) poorly structured sgRNAs were excluded; 3) cover approximately every 50 bp over the target sequences. First, candidate guides in rRNA (4324) and mtRNA (1907) were outputted by scanning all the protospacer adjacent motif (PAM) sequences. Meanwhile, the transcriptome of hg38 (excluded rRNA and mitochondrial RNA) was also scanned for all CRISPR sites as an off-target list. Second, we excluded off-target guides and filtered poorly structured guides, including a). G/C frequency too high (> 15/20) or too low (< 5/20); b). Homopolymer: more than 5 consecutive repeated nucleotides; c). Dinucleotide repeats: the same two nucleotides alternate for > 3 repeats; d). Hairpin: complementary subsequences near the start and end of a binding site, causing a hairpin. And we got 1191 rRNA guides and 1425 mtRNA guides as qualified guides. Next, we downloaded guides designed by Benchling to score our guides and remove redundant guides (overlapped with each other), and kept guides with higher on-target and off-target scores. Thus, we got a pool of filtered guides. Finally, we manually added some guides to cover the non-guide regions and SNP sites and obtained a final sgRNA pool containing 302 guides targeting rRNA sequences and 315 guides targeting mtRNA sequences. The DNA templates of final sgRNAs were synthesized through a one-step PCR using two paired primers to achieve the addition of T7 promoter and guide RNA scaffold sequences (Primers for preparation of sgRNA DNA templates were shown in **Supplementary Table 2**). The final sgRNAs were in vitro transcribed using T7 RNA polymerase (NEB, E2050) and stored at -80 °C.

### DETECTOR-seq library preparation

The 17.5 μl of remaining samples with similar cDNA content (ΔCt of ACTB-ee < 1) were pooled for 3-, 4- or 5-plex library preparation. The pooled cDNAs were pre-amplified using SeqAmp DNA Polymerase (Clontech, 638509) with the following PCR setup: initial denaturation at 94 °C for 1 min, denaturation at 98 °C for 15 s, annealing at 55 °C for 30 s, elongation at 68 °C for 30 s, and final elongation at 68 °C for 10 min. Pre-amplification was repeated for 6 cycles, and the DNA was cleaned and size-selected using Hieff NGS DNA selection Beads (Yeasen, 12601ES56) at a ratio of 1:0.8 of DNA to beads twice. The DNA was eluted with the 20 μl CRISPR-Cas9 reaction mix consisting of 300 ng sgRNAs for rRNA sequences, 40 ng sgRNAs for mtRNA sequences, 1× NEBuffer3.1 and 1 μM Cas9 Nuclease, *S. pyogenes* (NEB, M0386). The 20-µl reaction mixture was incubated at 37 °C for 60 min and heat-inactivated at 65°C for 5 min. Following the depletion of DNA fragments derived from rRNAs and mtRNAs, the remaining DNA samples with complete library structure were amplified for 16-18 cycles depending on the initial input. The final clean-up was conducted at a ratio of 1:1 of DNA to beads. The library concentration was measured using a Qubit dsDNA HS Assay Kit (Thermo Fisher, Q32854), and the size distribution of the library was assessed using an Agilent 2100 Bioanalyzer with a High-Sensitivity DNA analysis kit (Agilent, 5067-4626). DETECTOR-seq libraries were sequenced on an Illumina HiSeqX platform to a depth of 10 million 150 bp paired-end reads per sample. Primers for PCR amplification of DETECTOR-seq were shown in **Supplementary Table 4**.

### Sequencing data processing

Raw sequencing data were demultiplexed using sabre (https://github.com/najoshi/sabre) according to sample barcodes. UMI sequences were extracted using UMI-Tools [46]. Adapters were removed by cutadapt, and read pairs with an average quality score below 30 in either read were removed. The remaining read pairs were then mapped to ERCC’s spike-in sequences, NCBI’s UniVec sequences, human rRNA sequences, human mtRNA sequences, human genome (hg38), and circular RNA sequentially using STAR (version 2.5.3a_modified) [47]. The UMI-Tools package was used to remove duplicated reads caused by PCR amplification. Finally, a gene count matrix was generated using featureCounts v1.6.2 [48] with the GENCODE v38 annotation. Unmapped reads were classified using kraken2 [49] to obtain microbe (including bacterial, archaeal, and viral) genus abundance. Potential contaminations in genera were filtered before downstream analysis as in previously published work [19]. We summarized all the datasets for the development, validation, and application of DETECTOR-seq in **Supplementary Table 8**.

### Quality control (Sample filtering and Gene filtering)

To filter low-quality DETECTOR-seq datasets, the following quality control criteria were used **(Supplementary Table 7)**: (1) raw reads > 4 M; (2) clean reads (reads remained after trimming low-quality and adapter sequences) > 3.8 M; (3) ribosomal RNA reads < 20%; (4) mitochondrial RNA reads < 20%; (5) genome-aligned reads > 2 M; (6) de-duplicated RNA reads > 0.1 M. Next, we retained genes with TPM > 1 in at least 50% of samples or filtered genes by filterbyExpr of edgeR package [50].

### Differential expression and functional enrichment analysis

The count matrix of gene expression or microbe genus abundance was normalized using the trimmed mean of M-values (TMM) method in the edgeR package [50]. Differential expression analysis was conducted using a quasi-likelihood method with FDR<0.1 to identify RNAs showing a selective distribution in paired *Plasma* and *EV* samples and to identify differentially expressed genes (DEGs) (|log2fold-change|>1 and FDR<0.05) between cancer patients and normal controls. GSEA analysis of GO and KEGG pathway was carried out using clusterProfiler [51].

### Enrichment analysis of RBP binding motifs/sites

After identifying selective RNAs showing significantly differential abundance between *Plasma* and *EV* transcriptomes, we conducted an enrichment analysis of RBP (RNA binding protein) binding motifs/sites using MEME SEA [52]. We first created a gene-wise “RBP binding hotspot” sequence set by expanding annotated exon junction sites upstream and downstream by 20 nt and combined with 5’ UTR and 3’ UTR regions (GENCODE v38), as these regions were reported to be frequently bound by RBPs [53]. Background sequences were extracted from 500 random subsets of cfRNAs whose abundance showed no significant difference between *Plasma* and *EV* (FDR>0.1). Database files of RBP binding motifs/sites for enrichment analysis were annotated from our previous research [53]. Finally, top enriched RBPs (ranked by *E*-value) were annotated and summarized, and sequence logo images were created from POSTAR3 database [53] using WebLogo [54].

### Deconvolution of cell types of origin

We applied Nu-SVR to deconvolve the fractions of cell-type-specific RNAs based on Tabula Sapiens version 1.0 (TSP), a multiple-donor whole-body cell atlas spanning 24 tissues and organs as previously reported [23].

### Cancer classification

We normalized and scaled gene expression and genus abundance for evaluating the cancer-differentiating capacity of human and microbial features in *Plasma* and *EV* cfRNAs. All of the 61 *Plasma* and 52 *EV* DETECTOR-seq datasets passed QC were used, thus including as many cases as possible in the training and test sets. Most of the cancer samples were paired between *Plasma* and *EV* (CRC samples: *Plasma* 23, *EV* 19, 19 of them were paired; LC samples: *Plasma* 19, *EV* 19, 18 of them were paired; NC samples: *Plasma* 19, *EV* 14, 7 of them were paired). The data were trained and tested with bootstrapping sampling, which was randomly repeated 20 times. For human RNAs, a quasi-likelihood method was used for the differential expression analysis in each bootstrapping procedure. In Figure 7C, differentially expressed features with |log2fold-change|>1 and FDR<0.05 overlapped with RNAs that were enriched in *Plasma* or *EV* (defined in Figure 6A) were further used to fit a random forest classifier. In Figure 7D, we selected the top 200 features ranked by FDR in each bootstrapping procedure. For microbial RNAs, we selected all of the microbe genera with |log2fold-change|>1 and FDR<0.1 in each bootstrapping procedure. For the combination of human RNAs and microbial RNAs, we combined human gene expression and genus abundance and selected the top 200 features ranked by FDR. The area under the receiver operating characteristic curves (AUROC) was calculated from the final probability using the pROC [55] package in R.

### Cost estimation

The cost for cell-free RNA library preparation of DETECTOR-seq was determined using the sum of the price for each component used in our protocol. The price of SMARTer Stranded Total RNA-Seq Kit v2-Pico Input Mammalian (TaKaRa, 634413) was searched on the official website of TaKaRa for the estimation of SMARTer-seq. The cost of Phospho-RNA-seq was estimated using T4 polynucleotide kinase (NEB, M0201S) and TruSeq small RNA kit (Illumina, RS-200-0012). In the case of SILVER-seq, there was no publicly available step-by-step protocol, thus the cost of SILVER-seq was estimated by the Ovation SoLo RNA-Seq Kit (NuGEN, 0500-96). In all cases, the prices listed in **Supplementary Tables 6 and 7** included sales tax. Because the costs of SMARTer-seq, Phospho-RNA-seq, and SILVER-seq were estimated using commercial kits (including additional selling costs and profits), for a fair comparison, we determined the cost of DETECTOR-seq as twice the calculated price.

## Declarations

### Ethics approval and consent to participate

This study was approved by the institutional review board of Peking University First Hospital (2018-15). Informed consent was obtained from all patients.

### Consent for publication

Not applicable.

### Availability of data and materials

Data generated with DETECTOR-seq are available at the Gene Expression Omnibus under accession number GSE216561. For benchmarking, we used the following datasets: GSE126049 (Phospho-RNA-seq), GSE131512 (SILVER-seq), and GSE174302 (SMARTer-seq). *For editors and reviewers: the data can be downloaded from the GEO with a secure token: cnwpoiwufhunbqd*.

### Funding and Acknowledgments

This work is supported by National Natural Science Foundation of China (32170671, 8237061277 and 81902384), the Capital’s Funds for Health Improvement and Research (CFH 2022-2-4075), the National Key Research and Development Plan of China (2022ZD0117700), Tsinghua University Guoqiang Institute Grant (2021GQG1020), Tsinghua University Initiative Scientific Research Program of Precision Medicine (2022ZLA003), Tsinghua University Spring Breeze Fund (2021Z99CFY022). This study was also supported by Bayer Micro-funding, Bio-Computing Platform of Tsinghua University Branch of China National Center for Protein Sciences.

### Authors’ contributions

HKW, QZ, and ZJL conceived and designed the project; HKW developed DETECTOR-seq and generated the datasets; SZ and PYW collected the clinical samples; QZ and HKW conducted the analyses; all authors wrote and approved the manuscript.

