## Supplemental Files for "Depletion-assisted multiplexed cell-free RNA sequencing reveals distinct human and microbial signatures in plasma versus extracellular vesicles"

### Supporting Information

#### Supplementary Figures

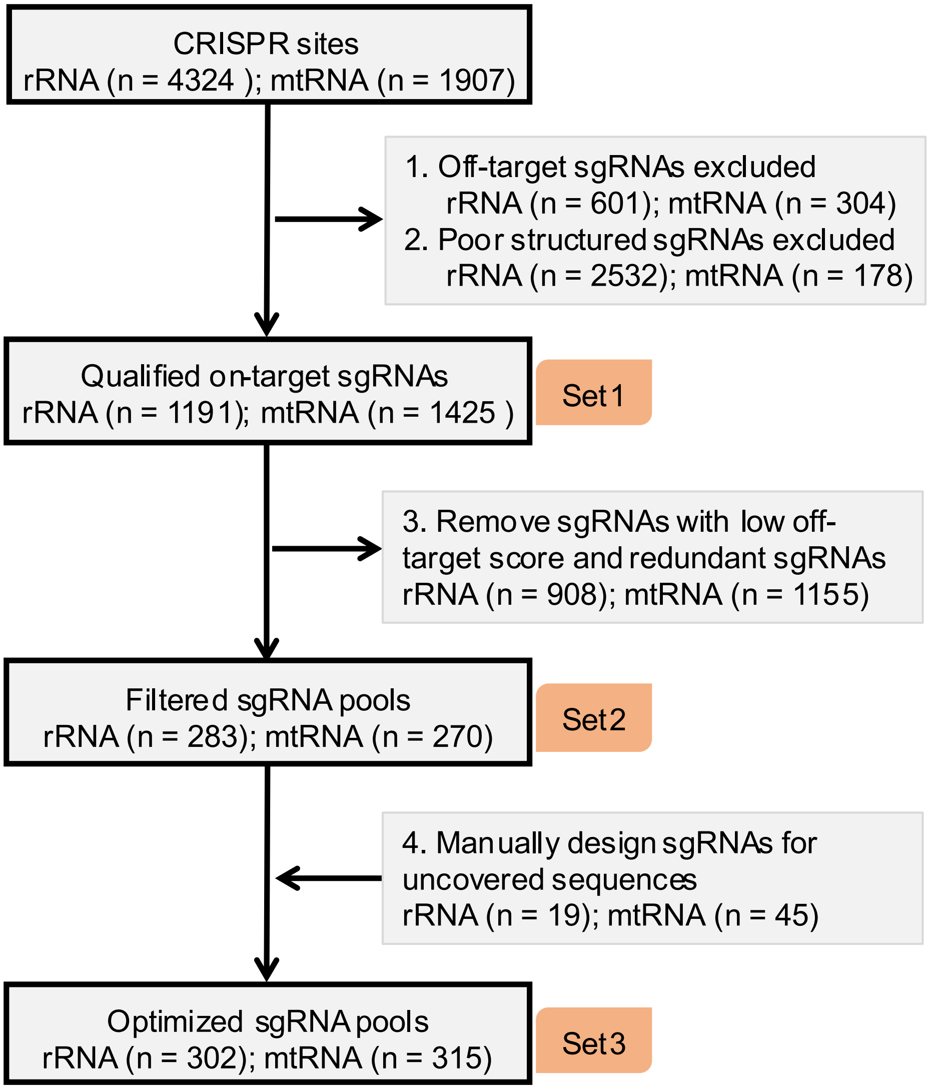

**Fig. S1. Design of sgRNAs.**

The workflow for the design of sgRNA pools for ribosomal RNAs (rRNAs) and mitochondrial RNAs (mtRNAs). See Methods for details.

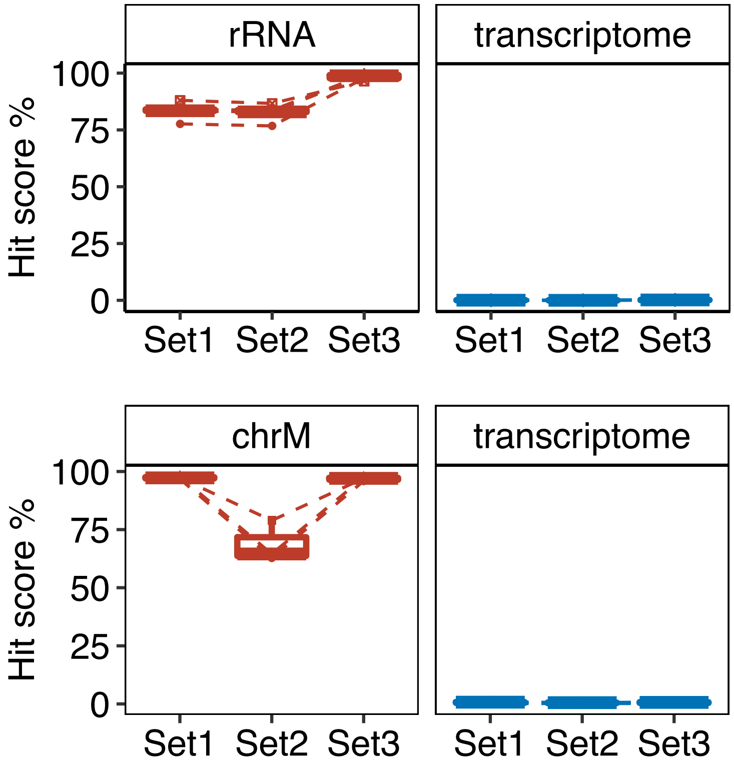

**Fig. S2. Evaluation of sgRNAs.**

In silico evaluation of hit scores for the sgRNA Set1, Set2, and Set3. Three plasma cfRNA datasets generated by DETECTOR-seq without the depletion of rRNA and mtRNA were used. chrM: RNAs derived from mitochondrial chromosome; transcriptome: plasma cell-free transcriptome eliminating rRNAs and mtRNAs.

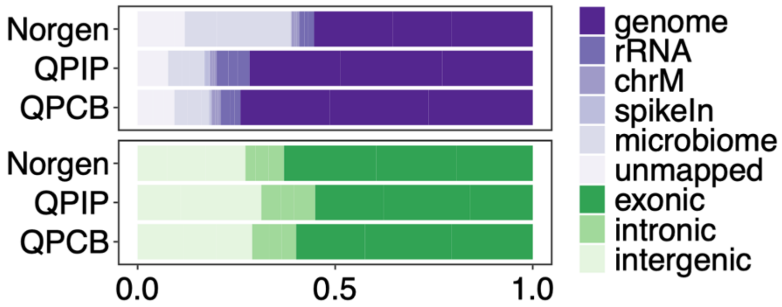

**Fig. S3. Comparison of plasma cell-free RNA extraction approaches.**

Read distributions of DETECTOR-seq datasets with three commonly used cell-free RNA extraction approaches (Norgen, QPIP, and QPCB). The top panel shows the average percentages of clean reads (reads remained after trimming low-quality and adapter sequences) assigned to human genome, rRNA, mtRNA, ERCC spikeIn, microbiome, and unmapped. The bottom panel shows the average percentages of genome-aligned reads mapped to exonic, intronic, and intergenic regions. Norgen: Plasma/Serum Circulating and Exosomal RNA Purification Kit; QPIP: QIAzol lysis, phenol-chloroform extraction, and isopropanol precipitation; QPCB: QIAzol lysis, phenol-chloroform extraction, and column binding.

A

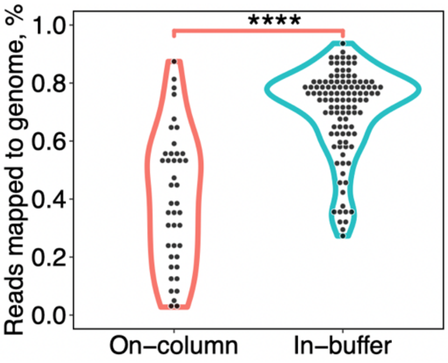

B

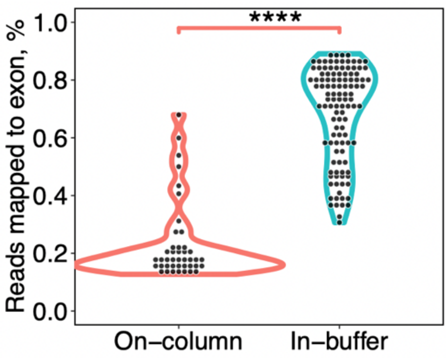

**Fig. S4. Comparison of residual cell-free DNA digestion approaches.**

(A) The percentages of clean reads aligned to human genome with different residual cell-free DNA digestion approaches. On-column: residual cell-free DNA was digested on the spin-column during RNA extraction; In-buffer: DNA was digested in aqueous buffer after RNA extraction. (B) The percentages of reads mapped to exons in the genome-aligned reads (N = 42 RNA samples using On-column approach and 113 RNA samples using In-buffer approach). ****: *P*-value < 0.0001, Wilcoxon rank sum test, two-tailed.

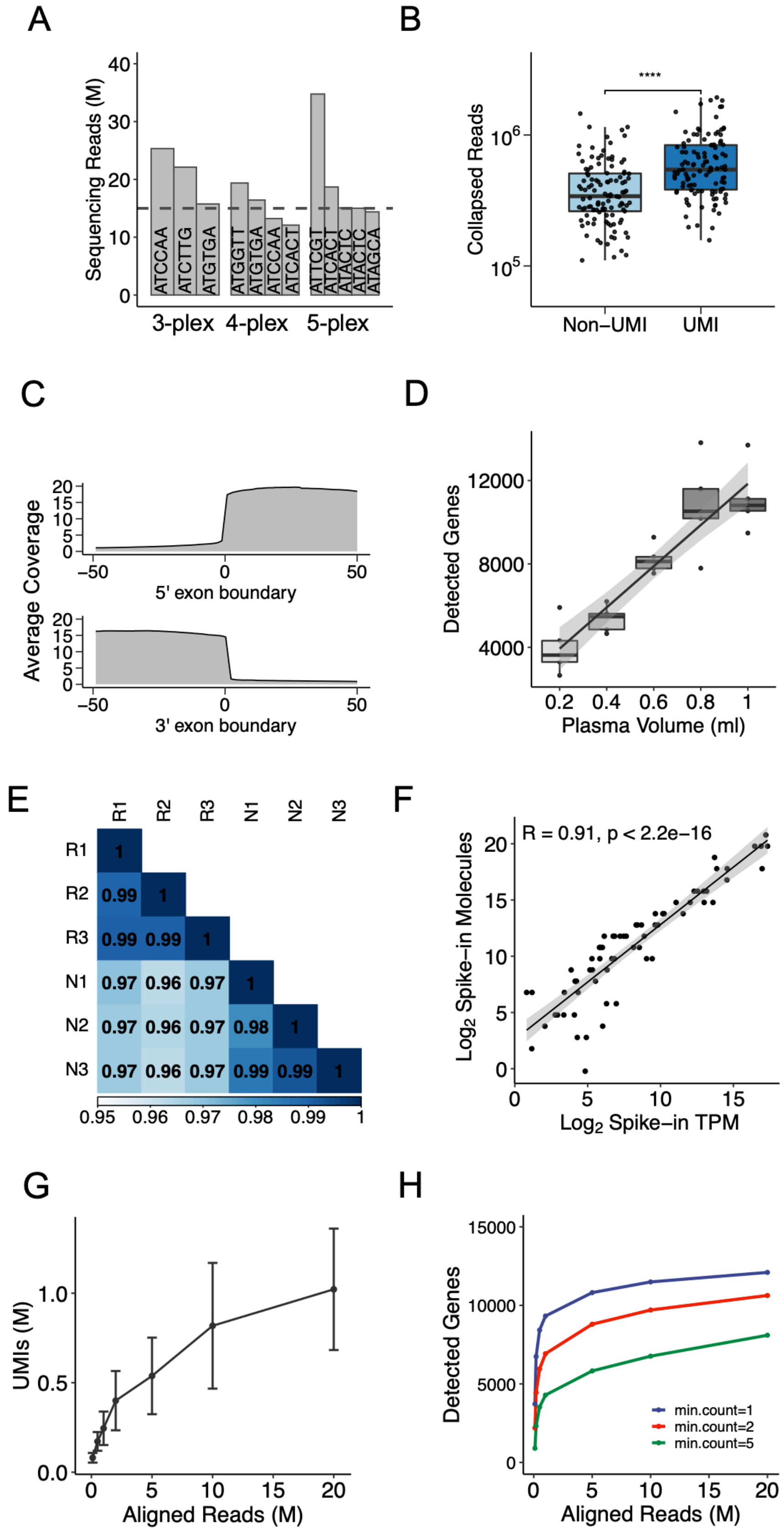
A
Sequencing Reads (M)
0
10
20
30
40
ATCCAA
ATCTTG
ATGTGA
ATGGTT
ATGTGA
ATCCAA
ATCACT
ATTCGT
ATCACT
ATACTC
ATACTC
ATAGCA
3-plex
4-plex
5-plex
B
****
Collapsed Reads
$10^6$
$10^5$
Non−UMI
UMI
C
Average Coverage
0
5
10
15
20
−50
0
50
5' exon boundary
3' exon boundary
D
Detected Genes
4000
8000
12000
0.2
0.4
0.6
0.8
1
Plasma Volume (ml)
E
R1
R2
R3
N1
N2
N3
0.95
0.96
0.97
0.98
0.99
1
F
R = 0.91, p < 2.2e−16
$Log_2$ Spike−in Molecules
$Log_2$ Spike−in TPM
0
5
10
15
20
G
UMIs (M)
0.0
0.5
1.0
0
5
10
15
20
Aligned Reads (M)
H
Detected Genes
0
5000
10000
15000
min.count=1
min.count=2
min.count=5
Aligned Reads (M)

#### Fig. S5. Analytical validation analysis of DETECTOR-seq's performance.

(A) Number of sequenced reads of each barcoded sample in each multiplexed library. The dashed line represents an expected number. (B) The number of collapsed reads with PCR duplicates removed by UMI or non-UMI methods. ****: P-value<0.0001, Wilcoxon rank sum test, two-tailed. (C) Average coverage across all the 5' and 3' exon boundary sites flanking upstream and downstream by 50 bp. (D) The number of detected genes in DETECTOR-seq libraries (n=5) with different input volumes of plasma. (E) Pearson correlation matrix of plasma samples from biological triplicates (N1–N3) and technical triplicates (R1–R3). (F) Pearson correlation between spike-in molecules and their reads sequenced by DETECTOR-seq for ERCC spike-in controls. (G) Numbers of detected UMIs and (H) detected genes (defined by three different minimum counts) at various subsampled genome-aligned read depths. The error bar represents the standard deviation of multiple samples (n=24). M: million.

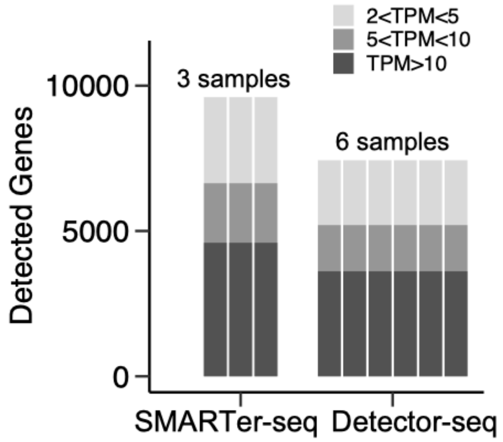

**Fig. S6. Cost-effectiveness of DETECTOR-seq compared to SMARTer-seq.**

The number of RNA samples can be processed for library preparation and detected genes for each sample. With a set budget of $300, SMARTer-seq could generate three cell-free RNA libraries and detect 9606 genes (TPM>2) for each library, while DETECTOR-seq could finish six cell-free RNA libraries and detect ~7433 genes (TPM>2) for each library. Although DETECTOR-seq detected fewer genes for each sample, it was much more cost-effective than SMARTer-seq because DETECTOR-seq could generate more libraries with the same budget.

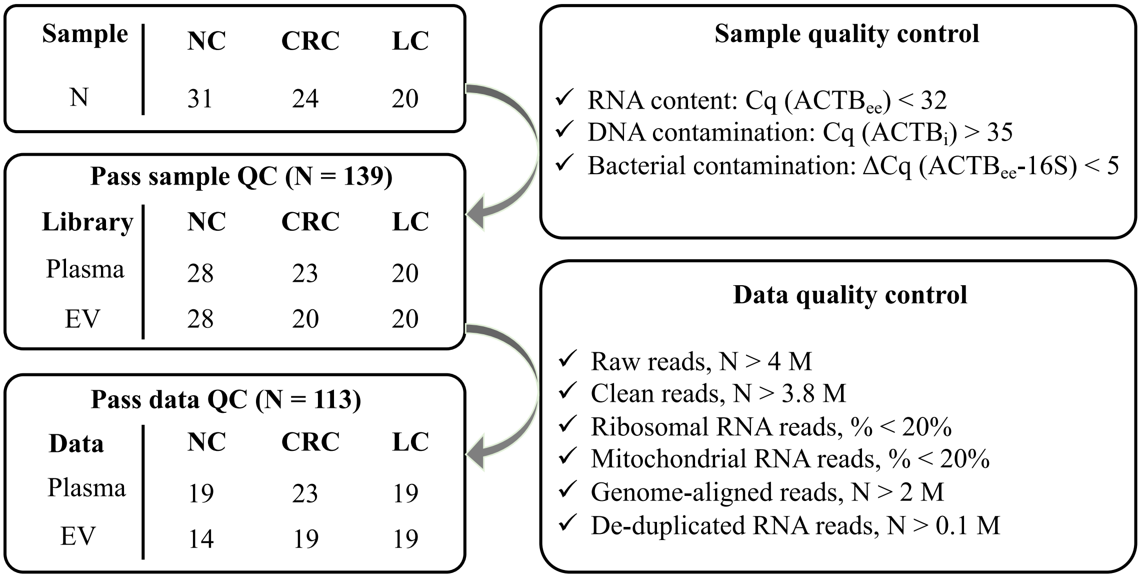

**Fig. S7. Summary of Quality control results for RNA samples and datasets.**

The left panel shows the number of samples, libraries, and datasets. And the right panel recapitulates the criteria of sample quality control and data quality control. NC: normal control; CRC: colorectal cancer; LC: lung cancer; $ACTB_{ee}$: the exon-exon junction region of *ACTB*; $ACTB_{i}$: the intron region of *ACTB*; 16S: the V4 region of 16S ribosomal RNA.

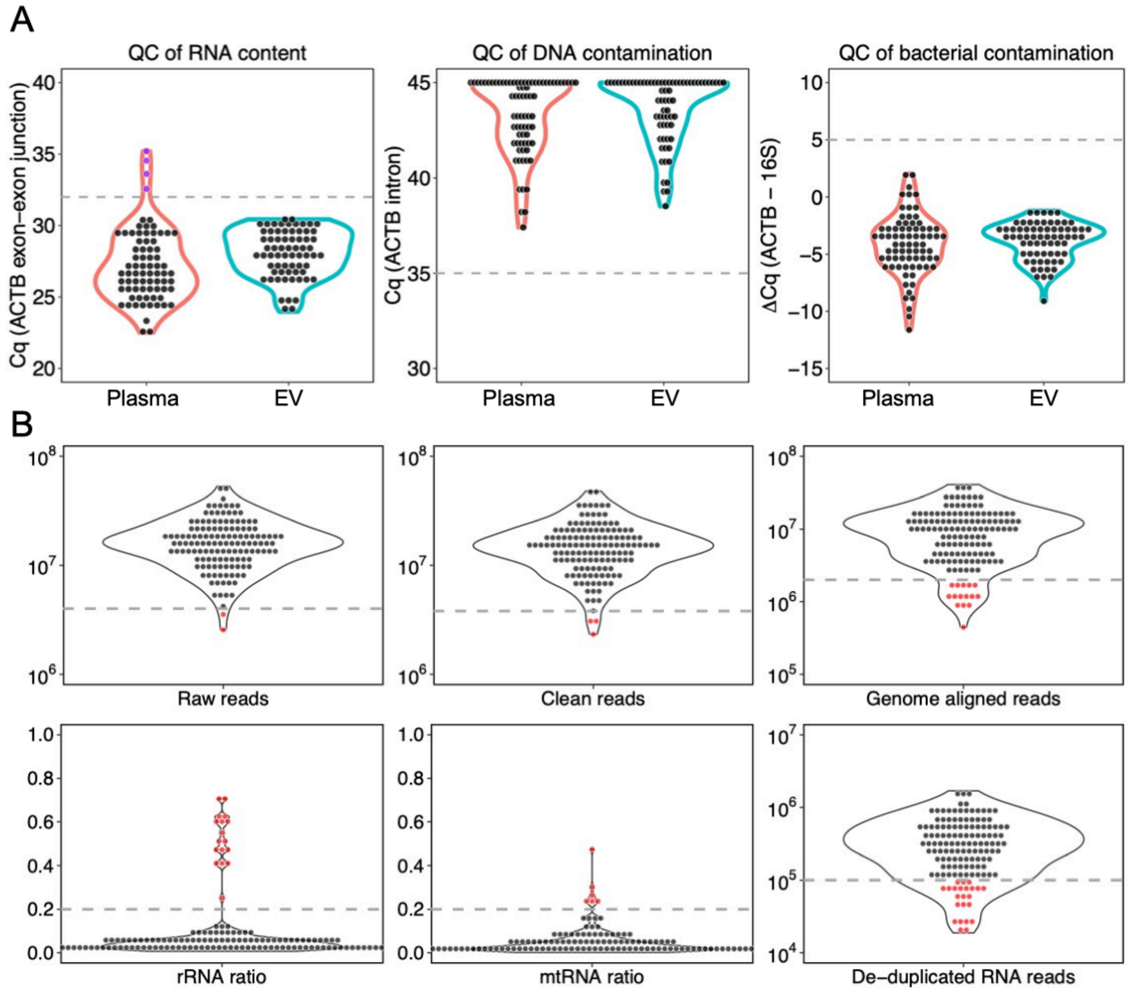

**Fig. S8. Quality control of Plasma and EV cfRNA samples and datasets.**

(A) Quality control results of RNA content, DNA contamination, and bacterial contamination. The estimation of RNA content was determined by the exon-exon junction region of *ACTB*. The level of residual DNA contamination was measured by amplifying a short fragment from *ACTB* within intron regions. We measured the bacterial contamination by the threshold cycle (Ct) value difference of the *ACTB* fragment spanning the exon-exon junction and a bacterial 16S ribosomal RNA V4 fragment. Primers were summarized in Table S1. Black dots represent RNA samples that passed the sample QC, while purple dots represent RNA samples that failed the sample QC. (B) The number of raw reads, clean reads, genome-aligned reads, and de-duplicated RNA reads as well as ribosomal RNA (rRNA) ratio and mitochondrial RNA (mtRNA) ratio were determined for each of the *Plasma* or extracellular vesicle (*EV*) RNA datasets generated by DETECTOR-seq. Black dots represent RNA datasets that passed the data QC, while red dots represent RNA datasets that failed the data QC. Dashed lines illustrate the cut-off values.

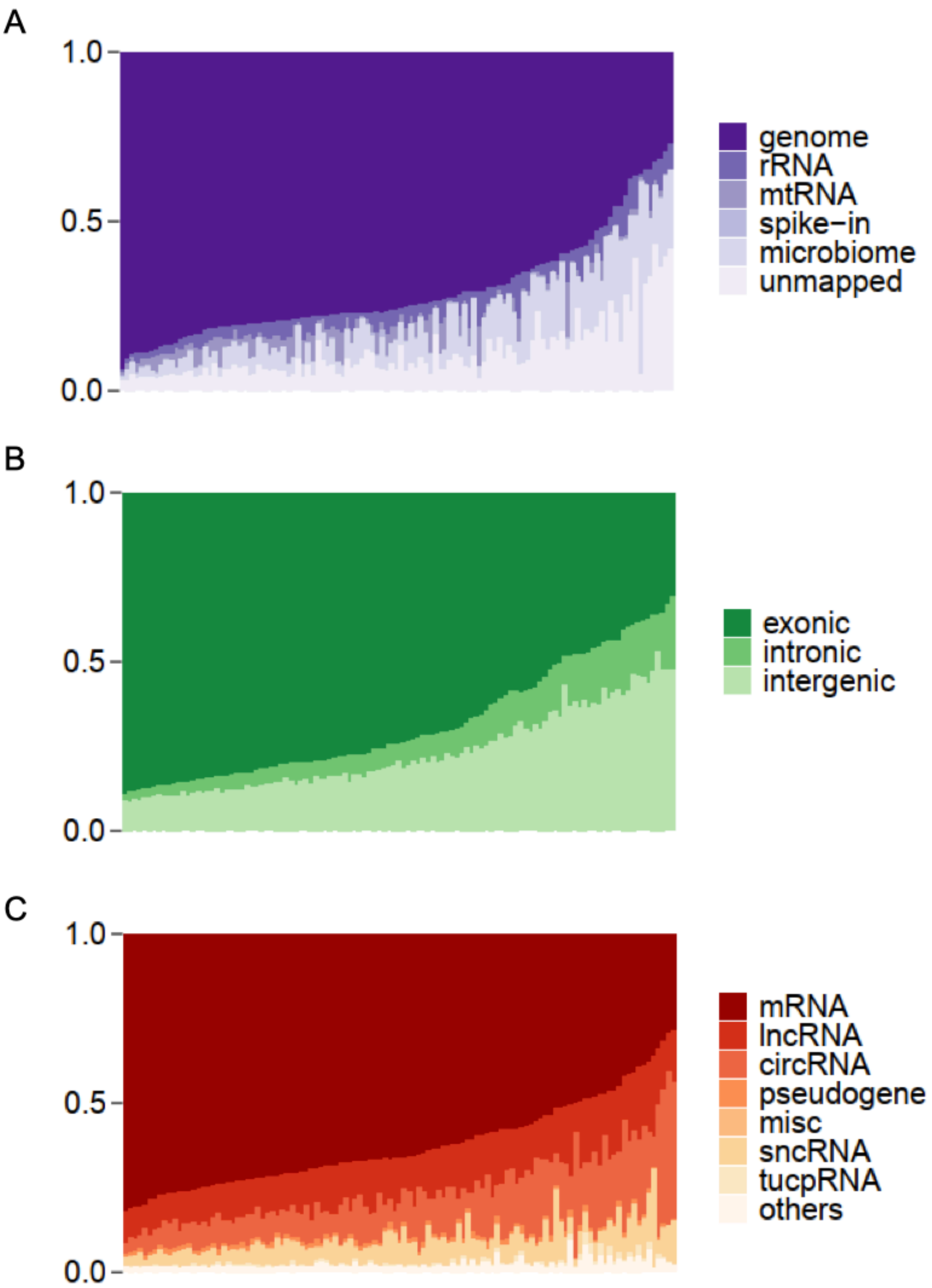

**Fig. S9. Read distributions across all samples that passed data quality control.**

(A) Percentages of clean reads assigned to human genome, rRNA, mtRNA, Spike-In, microbiome, and unmapped. (B) Percentages of genome-aligned reads derived from exonic, intronic, and intergenic fractions. (C) Percentages of exonic reads attributed to different RNA biotypes.

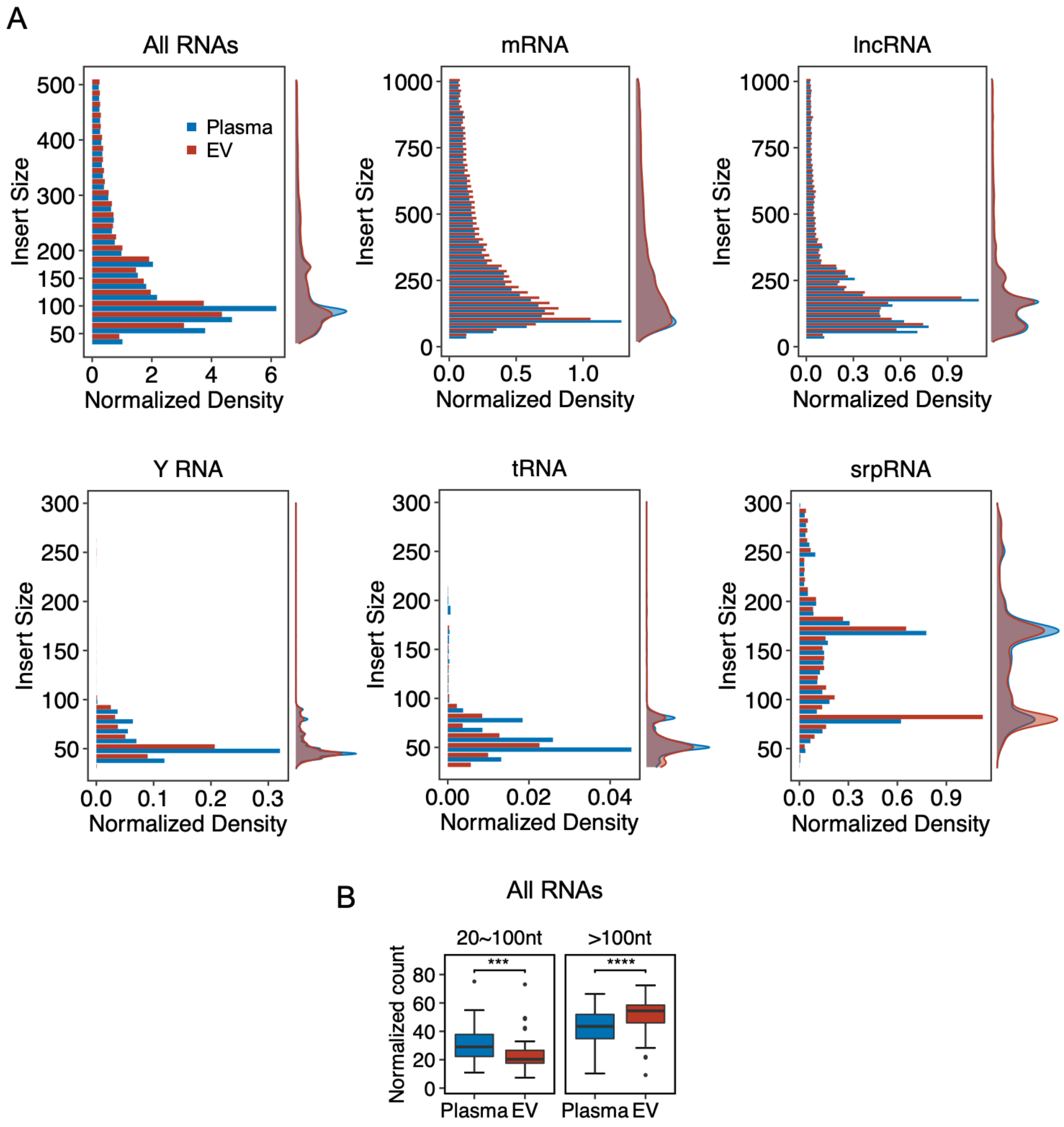

**Fig. S10. Fragment size of *Plasma* and *EV* cfRNAs.**

(A) The distributions of fragment size for various RNA species including mRNA, lncRNA, Y RNA, tRNA, and srpRNA in the cell-free transcriptomes of Plasma and EV. (B) The normalized count of 20~100nt group and >100nt group in Plasma and EV.

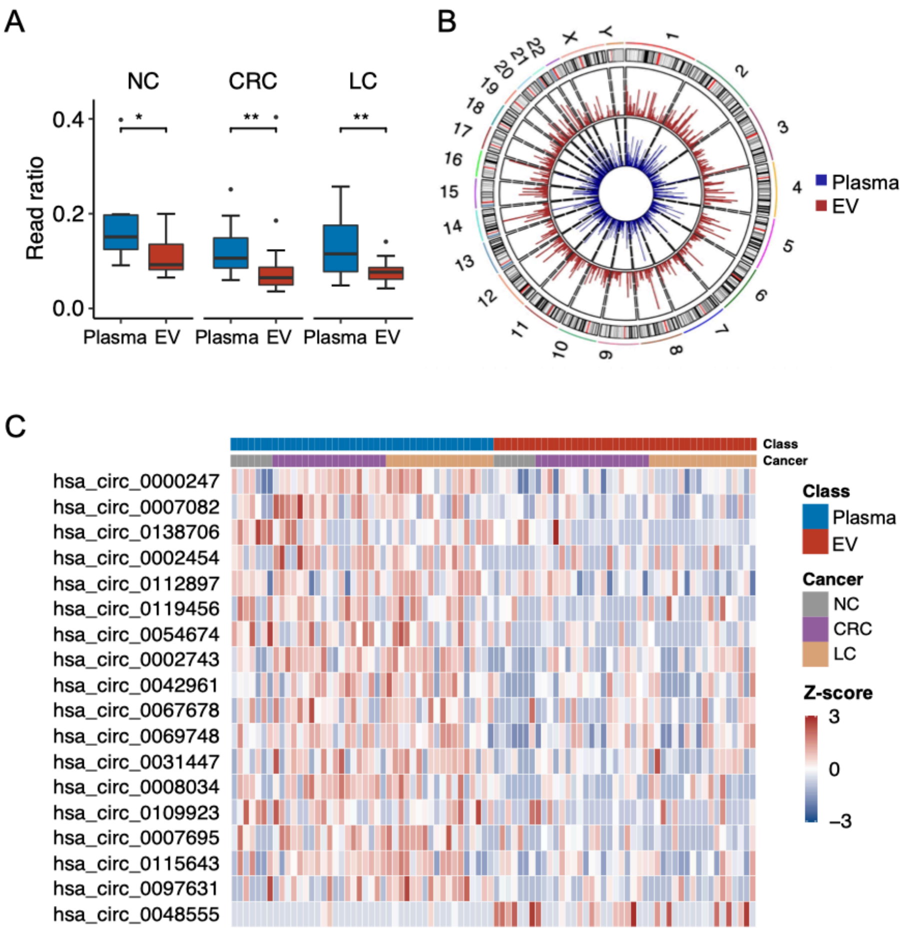

**Fig. S11. CircRNA profiles in *Plasma* and *EV*.**

(A) Read ratio of circRNAs in *Plasma* or extracellular vesicle (*EV*) cfRNAs across different kinds of samples. NC: normal control; CRC: colorectal cancer; LC: lung cancer. (B) Profiles of circRNAs across human chromosomes in *Plasma* and *EV* transcriptomes. (C) Identification of circRNAs that were enriched in Plasma and EV. Enriched RNAs were defined as differentially expressed RNAs between *Plasma* and *EV* with a cutoff of |Fold-change)|>1 and FDR<0.1.

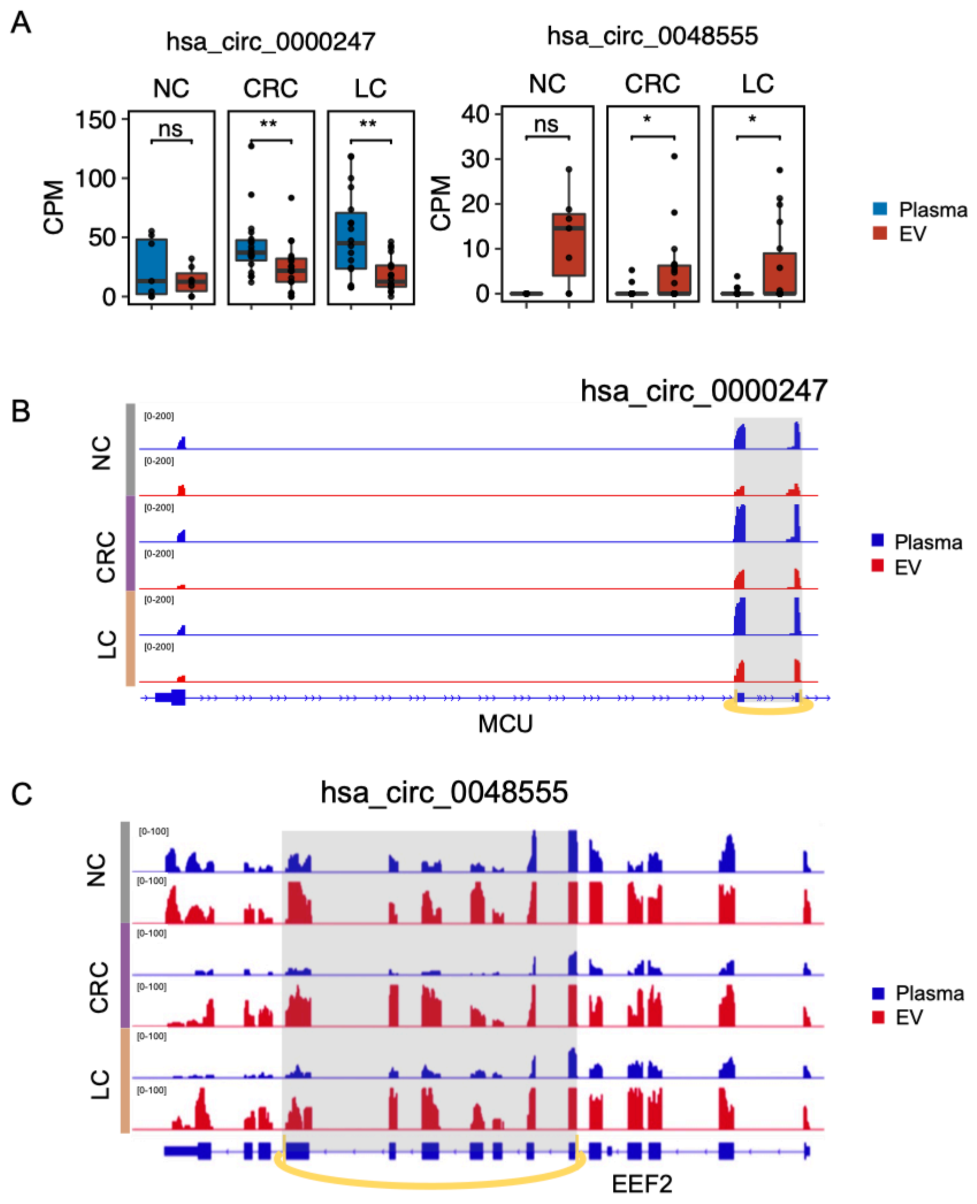

**Fig. S12. Representative circRNAs that were enriched in *Plasma* or *EV*.**

(A) Box plots showing the relative expression levels of hsa_circ_0000247 and hsa_circ_0048555 across different kinds of samples. (B) Read distributions of hsa_circ_0000247 in *Plasma* and *EV* show the differential expression pattern (in the gray box). (C) Read distributions of hsa_circ_0048555 in *Plasma* and *EV* show the differential expression pattern (in the gray box).

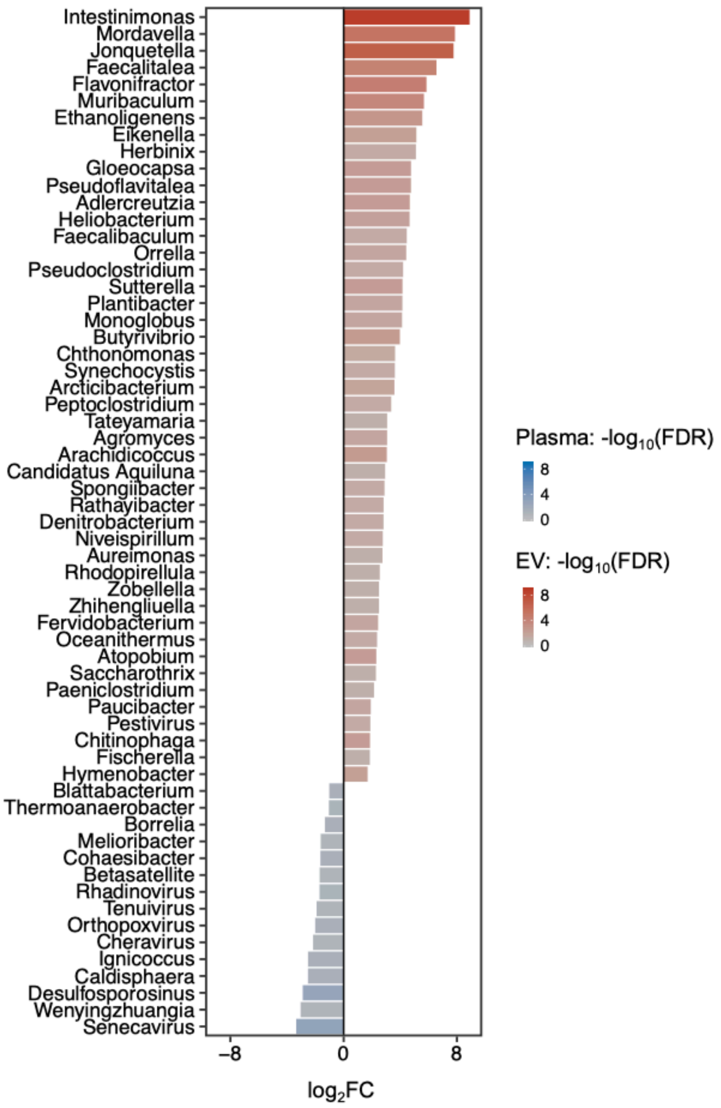

**Fig. S13. Microbe genera that were enriched in *Plasma* or *EV*.**

The microbe genera showed a significantly different abundance between *Plasma* and *EV* with a cutoff of $|\log_2$fold-change$|>1$ and FDR<0.1. FC: fold-change; FDR: false discovery rate.

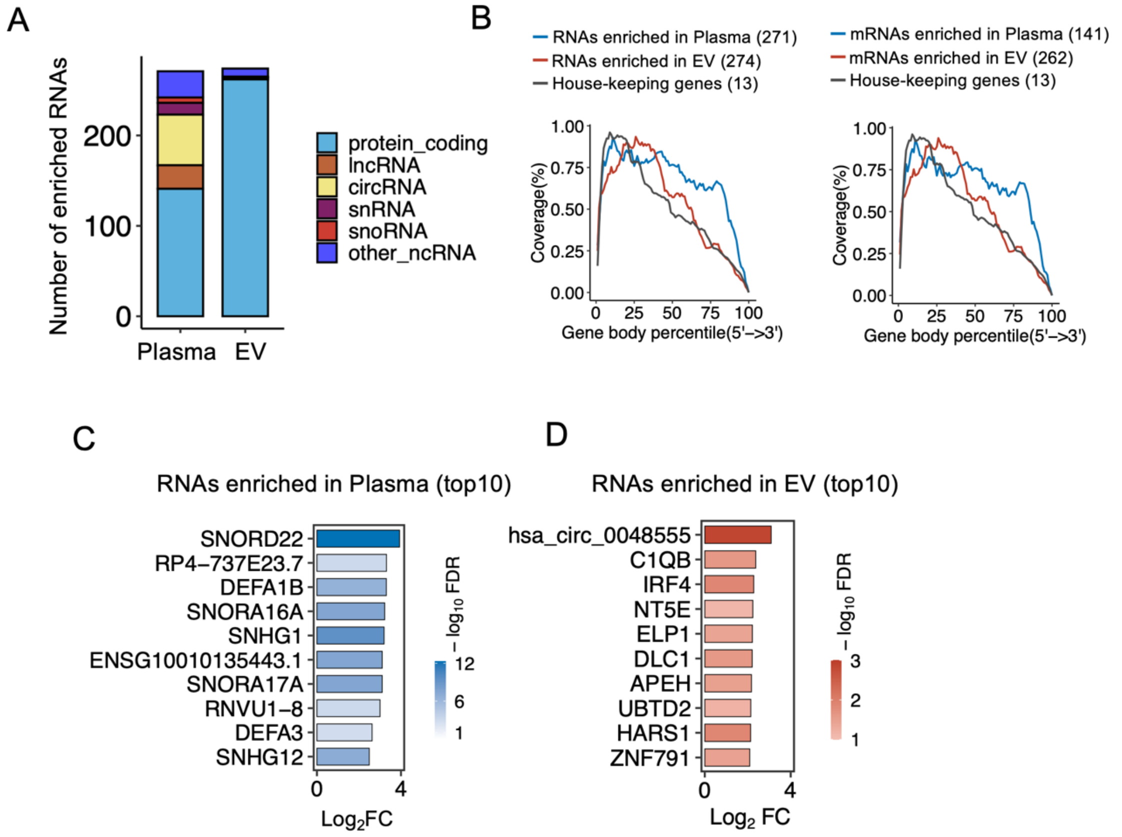

**Fig. S14. Enriched RNAs in *Plasma* or *EV*.**

(A) The number of RNAs that were enriched in *Plasma* or *EV* counted by RNA species. (B) Gene body coverage, computed as an average across RNAs(left) or protein coding genes(right) enriched in *Plasma*/*EV* and house-keeping genes. RNAs that were enriched in *Plasma* exhibited a higher gene body coverage than RNAs that were enriched in *EV*, indicating its degradation-resistant nature. *Plasma*: n = 44 samples; *EV*: n = 44 samples. C Bar plots showing the top 10 RNAs that were enriched in *Plasma*. (D) Bar plots showing the top 10 RNAs that were enriched in *EV*.

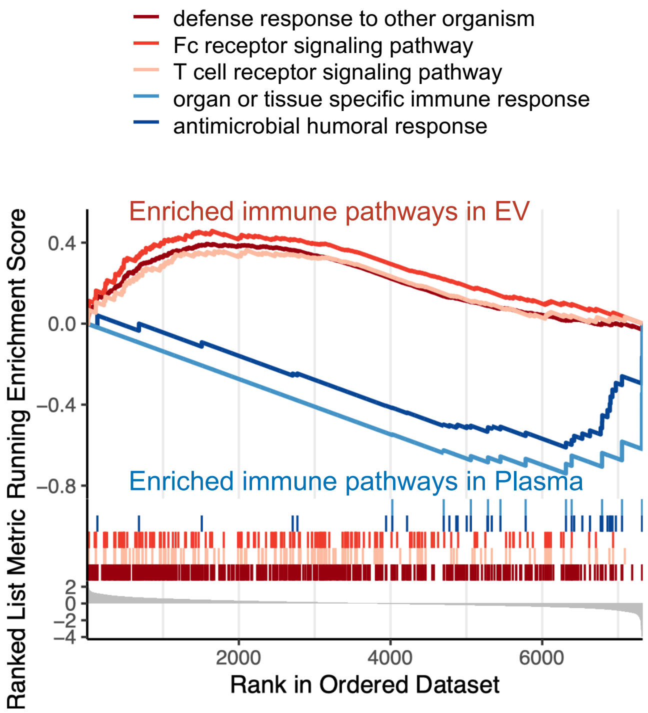

**Fig. S15. Enriched immune pathways in *Plasma* or *EV*.**

Gene set enrichment analysis exhibiting distinct immune response pathways associated with cell-free RNAs that were enriched in *Plasma* or *EV*.

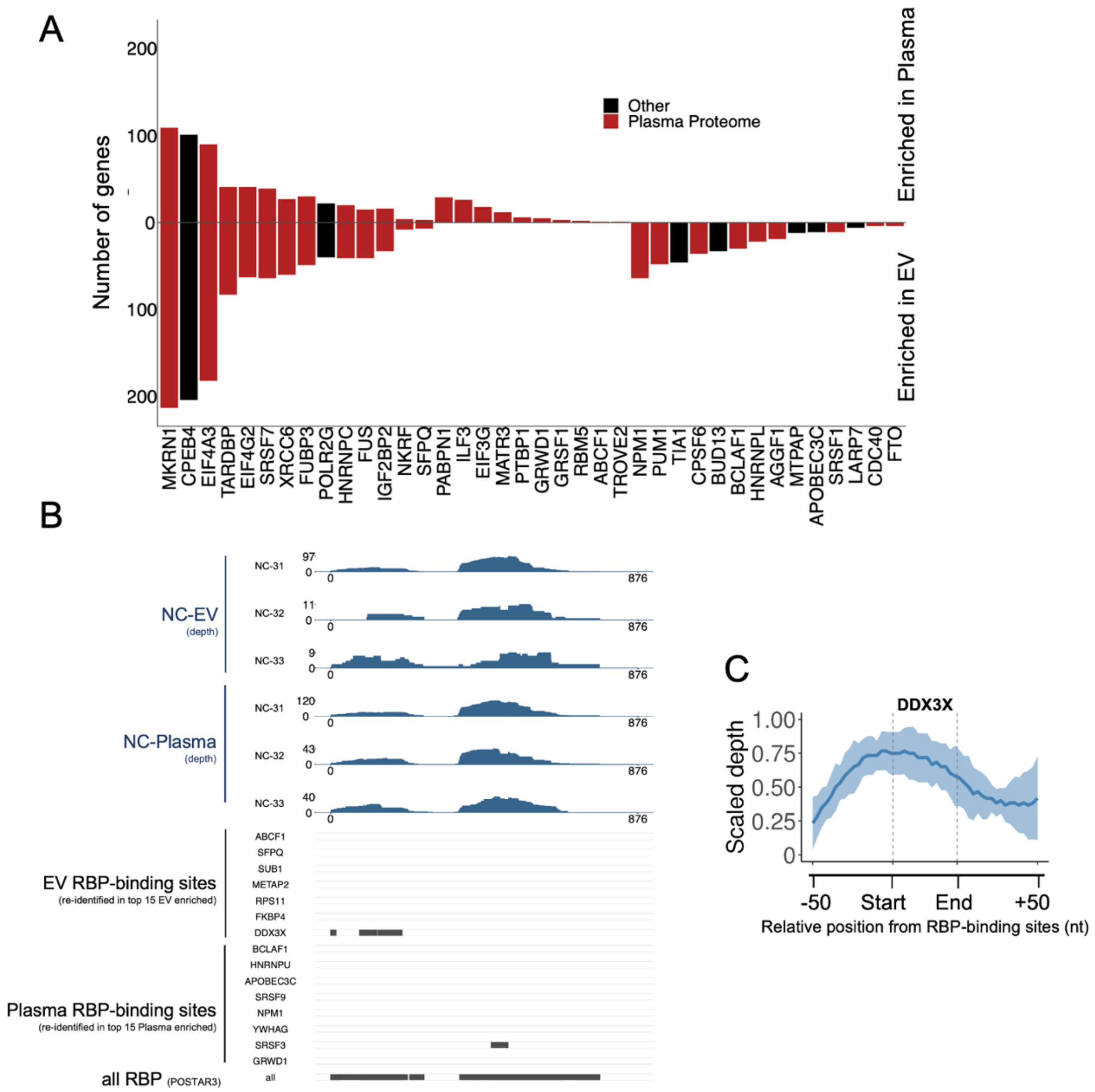

**Fig. S16. The predicted prevalent RBPs in the selective RNAs enriched in *Plasma* or *EV*.**

(A) The occurrences of RBP binding motifs/sites were scanned with MEME FIMO **(1)** using the "RBP binding hotspot" annotated in POSTAR3 **(2)**. Results were further filtered by q value < 0.05. Here, the predicted RBPs in the selective cfRNAs enriched in *Plasma* or *EV* were ranked by gene count, with RBPs reported in plasma proteome **(3)** shown in red. RBP: RNA-binding protein. (B) RNA peaks near EV or Plasma enriched RBP-binding sites (C) Coverage around annotated RBP-binding sites and the region of binding sites show higher coverage of cfRNA(<65 nt, similar to most CLIP-seq) fragments.

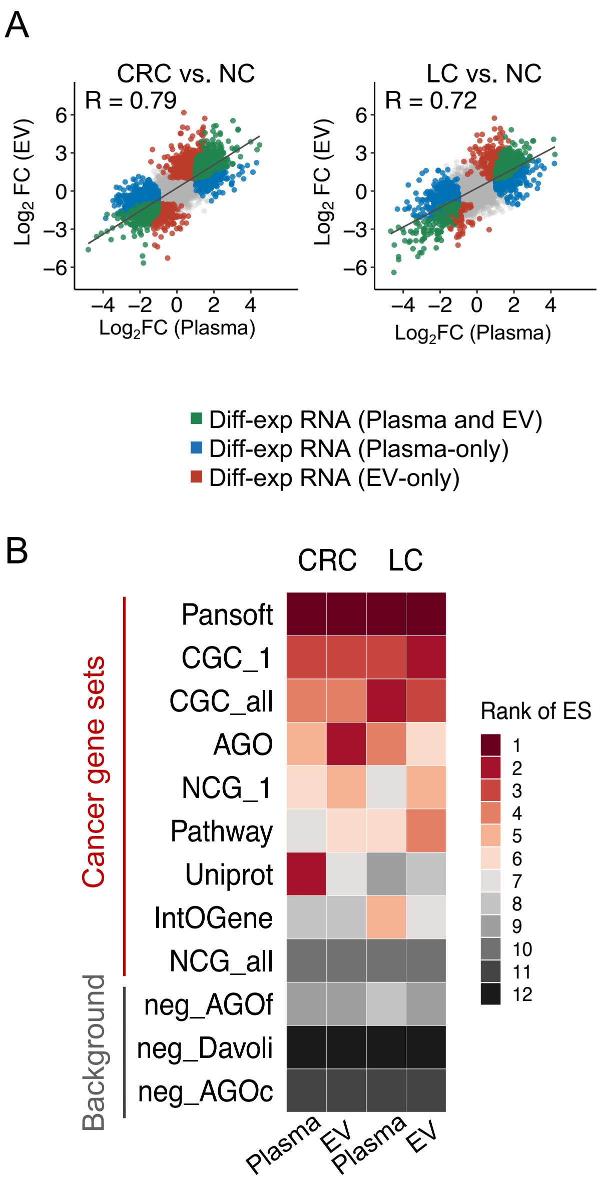

**Fig. S17. Cancer-relevant RNAs detected in *Plasma* and *EV*.**

(A) Common and specific cancer-relevant RNAs (DEGs, differentially expressed genes identified between cancer patients versus normal controls) detected in *Plasma* and *EV*. The fold-change of these DEGs in *Plasma* and *EV* showed a high consistency, indicating that *Plasma* and *EV* cfRNAs have similar potential for cancer detection. Pearson correlation coefficient (R) of the regression line was calculated. EV: extracellular vesicle; NC: normal control; CRC: colorectal cancer; LC: lung cancer. (B) To determine if these DEGs enriched cancer-related signals, we curated known cancer gene sets as well as non-cancer gene sets (as background) and calculated an enrichment score defined as the overlap of DEGs with certain gene sets normalized by the size of the gene sets. As expected, the enrichment scores of cancer gene sets (8/9, except NCG_all) were consistently elevated compared to those of three non-cancer gene sets. These results indicate that DETECTOR-seq captures the cancer signals both in *Plasma* and *EV* cfRNAs. We collected the cancer gene sets from the following resources: Tier 1 from COSMIC Cancer Gene Census **(4)** version 92 where genes must possess a documented activity relevant to cancer with evidence (CGC_1), Tier 1 and Tier 2 of CGC which included genes with strong indications of a role in cancer but with less extensive available evidence (CGC_all), canonical cancer drivers of the Network of Cancer Genes 6 (NCG_1), canonical cancer drivers and candidate cancer drivers of NCG (NCG_all), the Atlas of Genetics and Cytogenetics in Oncology and Hematology (AGO), UniprotKB (Uniprot), PanSoftware (PanSoft) **(5)**, Integrative OncoGenomics (IntOGen) **(6)**, pathway in cancer of KEGG (Pathway). Meanwhile, we collected the gene sets unrelated to cancers as negative controls from the following resources: a conservative version of the negative AGO list (neg_AGOc), a list derived from AGO (neg_AGOf), and a list of known non-driver genes (neg_Davoli) **(7)**.

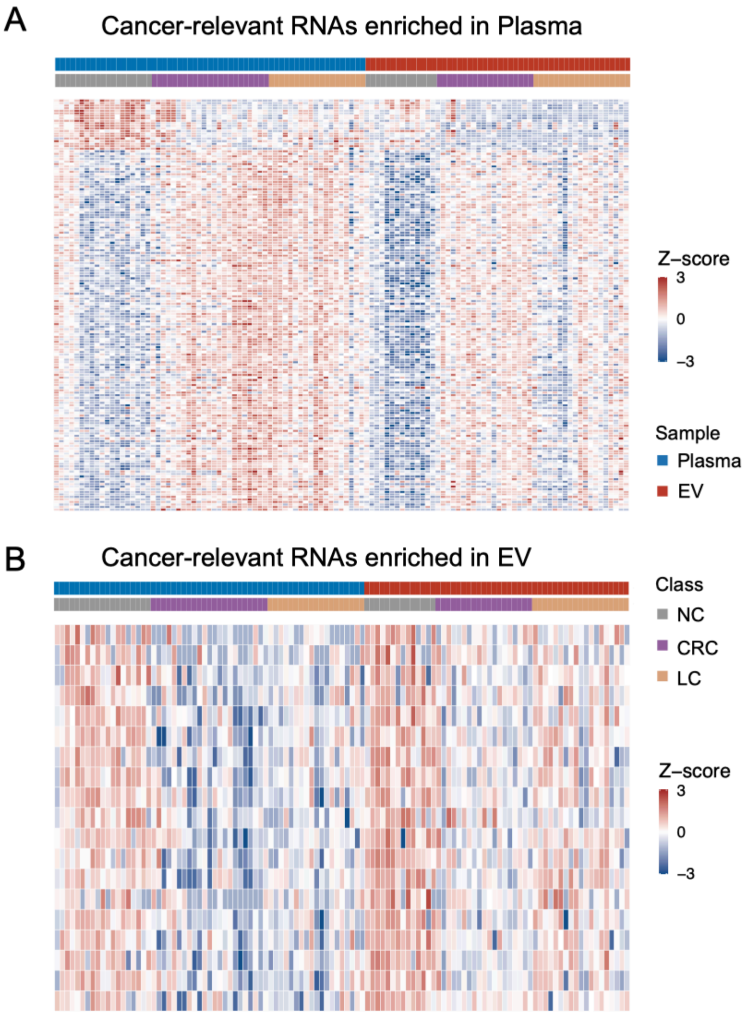

**Fig. S18. The relative abundance levels of cancer-relevant cell-free RNAs enriched in *Plasma* or *EV*.**

(A) Heatmap showing the expression patterns of cancer-relevant cell-free RNAs that were enriched in *Plasma*. (B) Heatmap showing the expression patterns of cancer-relevant cell-free RNAs that were enriched in *EV*. Cancer: CRC and LC were collectively labeled as cancer. Cancer-relevant RNAs were defined as differentially expressed RNAs ($|\log_2$(fold-change)$|>1$ and FDR<0.05) between cancer patients and normal controls. Selective RNAs in *Plasma* or *EV* were defined as differentially expressed RNAs (Fold-change|>1 and FDR<0.1) between *Plasma* and *EV*.

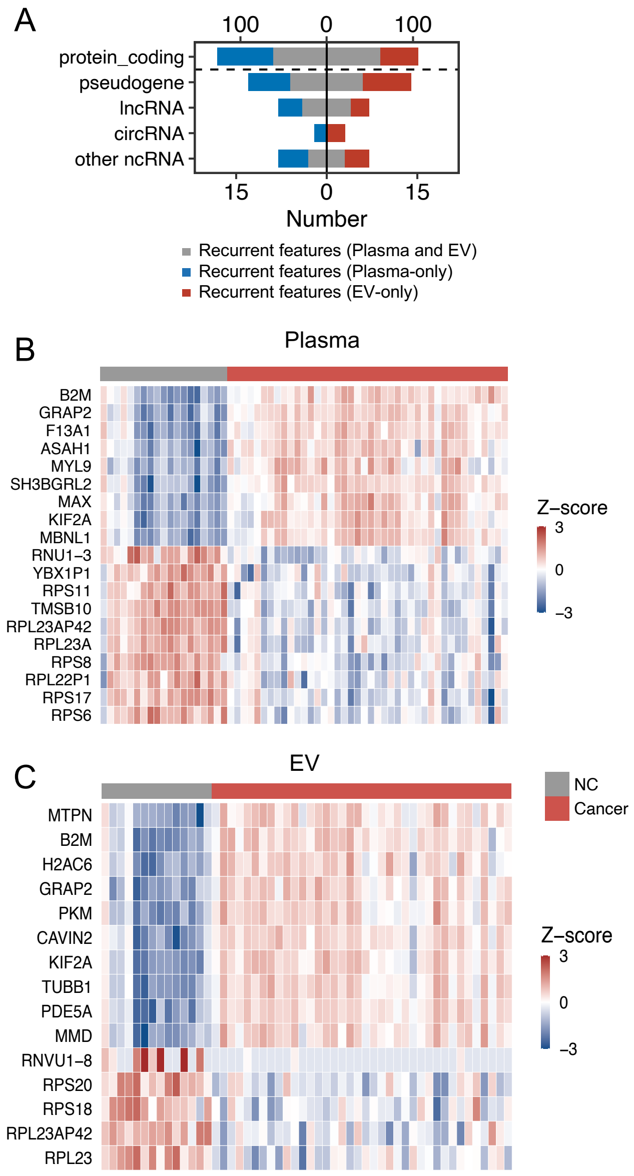

**Fig. S19. The expression patterns of recurrent cancer-relevant cell-free RNAs.**

(A) Bar plots showing the common and distinct recurrent cell-free RNA features in *Plasma* and *EV* counted by RNA species. (B) Heatmap showing the expression patterns of recurrent cancer-relevant cell-free RNAs in *Plasma*. (C) Heatmap showing the expression patterns of recurrent cancer-relevant cell-free RNAs in *EV*. Recurrent RNA features were identified as the top 200 differentially expressed RNAs between cancer patients and normal controls in all of the 20 bootstrap samplings.

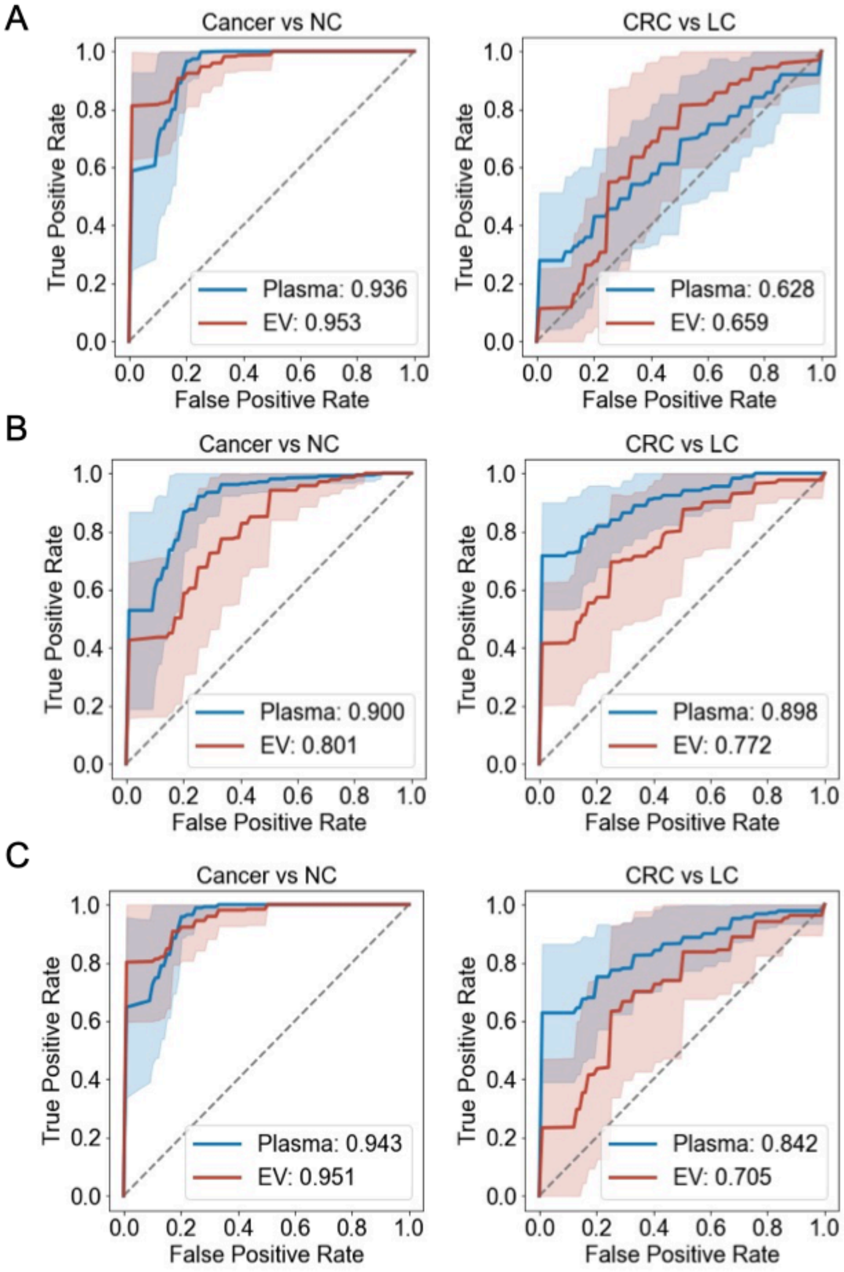

**Fig. S20. The performances of cell-free RNAs for cancer classification.**

(A) The performance of human-derived cell-free RNAs for cancer classification. (B) The performance of microbe-derived cell-free RNAs for cancer classification. (C) The performance of combination of human and microbial RNAs for cancer classification. EV: extracellular vesicle; CRC: colorectal cancer; LC: lung cancer; Cancer: CRC and LC.

#### Supplementary Tables

##### Table S1. Quality control primers for *Plasma* and *EV* RNA samples.

| Gene | Region | Primers | |
|---|---|---|---|
| ACTB | exon-exon junction | Forward | CTGAACCCCAAGGCCAACC |
| ACTB | exon-exon junction | Reverse | GAGGCGTACAGGGATAGCAC |
| ACTB | intron | Forward | TAAAGCGGCCTTGGAGTGTG |
| ACTB | intron | Reverse | GAACACGGCTAAGTGTGCTG |
| 16S rRNA | V4 | Forward | GTGCCAGCMGCCGCGGTAA(8) |
| 16S rRNA | V4 | Reverse | GGACTACHVGGGTWTCTAAT(8) |

**Table S2. Primers for the preparation of sgRNA DNA templates.**

| Primer name | Primer sequence |
|---|---|
| sgRNA Pool Forward | TAATACGACTCACTATAGGNNNNNNNNNNNNNNNNNN NNGTTTTAGAGCTAGAAATAGC |
| sgRNA Pool Reverse | TCGGTGCCACTTTTTCAAGTTGATAACGGACTAGCCTT ATTTTAACTTGCTATTTCTAGCTCTAAAAC |
| 5' Universal | ACGATCGTCGACAGCAAGCTAATACGACTCACTATAGG |
| 3' Universal | AAAAGCACCGACTCGGTGCCACTTTTTCAAGTTGATA |

This table shows primers for the preparation of sgRNA DNA templates. T7 promoter and sgRNA scaffold sequences were added through one-step PCR using two paired primers shown in the table. The target-specific region of sgRNAs was marked as 20 random nucleotides (N).

**Table S3. Primers for the reverse transcription of DETECTOR-seq.**

| Primer name | Primer sequence |
|---|---|
| TSO-UMI | Biotin-GTGTGCTCTTCCGATCTCGNNNNNNNNrGrGrG |
| N6-barcode1 | Biotin-ACGACGCTCTTCCGATCTAT**GGTT**NNNNNN |
| N6-barcode2 | Biotin-ACGACGCTCTTCCGATCTAT**TCGT**NNNNNN |
| N6-barcode3 | Biotin-ACGACGCTCTTCCGATCTAT**CACT**NNNNNN |
| N6-barcode4 | Biotin-ACGACGCTCTTCCGATCTAT**ACTC**NNNNNN |
| N6-barcode5 | Biotin-ACGACGCTCTTCCGATCTAT**AGCA**NNNNNN |
| N6-barcode6 | Biotin-ACGACGCTCTTCCGATCTAT**TTCC**NNNNNN |
| N6-barcode7 | Biotin-ACGACGCTCTTCCGATCTAT**GTGA**NNNNNN |
| N6-barcode8 | Biotin-ACGACGCTCTTCCGATCTAT**CTTG**NNNNNN |
| N6-barcode9 | Biotin-ACGACGCTCTTCCGATCTAT**CCAA**NNNNNN |
| N6-barcode10 | Biotin-ACGACGCTCTTCCGATCTAT**AAGG**NNNNNN |
| N6-barcode11 | Biotin-ACGACGCTCTTCCGATCTAT**GAAC**NNNNNN |
| N6-barcode12 | Biotin-ACGACGCTCTTCCGATCTAT**TGAG**NNNNNN |

Nucleotides marked in bold indicate the barcode sequences.

**Table S4. Primers for the PCR amplification of DETECTOR-seq.**

| Primer name | Primer sequence |
|---|---|
| 5'1 PCR1 | Biotin-AGAAGACGGCATACGAGAT**CGAGTAAT**GT GACTGGAGTTCAGACGTGTGCTCTTCCGATCT*C |
| 3'1 PCR1 | Biotin-CGACCACCGAGATCTACAC**TATAGCCT**AC ACTCTTTCCCTACACGACGCTCTTCCGATCT*A |
| 5'2 PCR1 | Biotin-AGAAGACGGCATACGAGAT**TCTCCGGA**GT GACTGGAGTTCAGACGTGTGCTCTTCCGATCT*C |
| 3'2 PCR1 | Biotin-CGACCACCGAGATCTACAC**ATAGAGGC**AC ACTCTTTCCCTACACGACGCTCTTCCGATCT*A |
| 5'3 PCR1 | Biotin-AGAAGACGGCATACGAGAT**AATGAGCG**GT GACTGGAGTTCAGACGTGTGCTCTTCCGATCT*C |
| 3'3 PCR1 | Biotin-CGACCACCGAGATCTACAC**CCTATCCT**AC ACTCTTTCCCTACACGACGCTCTTCCGATCT*A |
| 5'4 PCR1 | Biotin-AGAAGACGGCATACGAGAT**GGAATCTC**GT GACTGGAGTTCAGACGTGTGCTCTTCCGATCT*C |
| 3'4 PCR1 | Biotin-CGACCACCGAGATCTACAC**GGCTCTGA**AC ACTCTTTCCCTACACGACGCTCTTCCGATCT*A |
| 5'5 PCR1 | Biotin-AGAAGACGGCATACGAGAT**TTCTGAAT**GT GACTGGAGTTCAGACGTGTGCTCTTCCGATCT*C |
| 3'5 PCR1 | Biotin-CGACCACCGAGATCTACAC**AGGCGAAG**AC ACTCTTTCCCTACACGACGCTCTTCCGATCT*A |
| 5'6 PCR1 | Biotin-AGAAGACGGCATACGAGAT**ACGAATTC**GT GACTGGAGTTCAGACGTGTGCTCTTCCGATCT*C |
| 3'6 PCR1 | Biotin-CGACCACCGAGATCTACAC**TAATCTTA**AC ACTCTTTCCCTACACGACGCTCTTCCGATCT*A |
| 5'7 PCR1 | Biotin-AGAAGACGGCATACGAGAT**AGCTTCAG**GT GACTGGAGTTCAGACGTGTGCTCTTCCGATCT*C |
| 3'7 PCR1 | Biotin-CGACCACCGAGATCTACAC**CAGGACGT**AC ACTCTTTCCCTACACGACGCTCTTCCGATCT*A |
| 5'8 PCR1 | Biotin-AGAAGACGGCATACGAGAT**GCGCATTA**GT GACTGGAGTTCAGACGTGTGCTCTTCCGATCT*C |
| 3'8 PCR1 | Biotin-CGACCACCGAGATCTACAC**GTACTGAC**AC ACTCTTTCCCTACACGACGCTCTTCCGATCT*A |
| 5' PCR2 | CAAGCAGAAGACGGCATACGAGA*T |
| 3' PCR2 | AATGATACGGCGACCACCGAGATCTACA*C |

Nucleotides marked in bold indicate the index sequences. *: phosphorothioate.

**Table S5. Cell-free RNA sequencing datasets for the comparison analysis.**

| Methods | Accession ID | Specimen | N samples | Reference |
|---|---|---|---|---|
| Phospho-RNA-seq | GSE126049 | Plasma | 15 | (9) |
| SILVER-seq | GSE131512 | Serum | 128 | (10) |
| SMARTer-seq | GSE174302 | Plasma | 373 | (11) |
| DETECTOR-seq | - | Plasma | 113 | - |

Datasets generated by Phospho-RNA-seq, SILVER-seq, and SMARTer-seq were downloaded from GEO (Gene Expression Omnibus) database.

**Table S6. The estimated per sample cost of library preparation for DETECTOR-seq.**

| Primers / Reagents | Manufacturer | Catalog # | Total units | Units per sample | Total samples | $ Total | $ Cost per sample |
|---|---|---|---|---|---|---|---|
| N6 barcode primer | Sangon | NA | NA | NA | 200 | 29 | 0.14 |
| SMARTScribe Reverse Transcriptase | Clontech | 639538 | 40000 | 200 | 200 | 1099 | 5.50 |
| dNTP Mixture | Takara | 4019 | 250 µl | 2 µl | 125 | 11 | 0.09 |
| PEG8000 | Beyotime | R0056-2ml | 2000 µl | 2 µl | 1000 | 9 | 0.01 |
| Template switch oligo | Sangon | NA | NA | NA | 2000 | 65 | 0.03 |
| Recombinant RNase Inhibitor | Takara | 2313A | 5000 | 20 | 250 | 64 | 0.25 |
| FastFire qPCR PreMix (SYBR Green) | TIANGEN | FP207-03 | 5000 rxn | 10 rxn | 500 | 1260 | 2.52 |
| SeqAmp DNA Polymerase | Clontech | 638509 | 250 | 2.5 | 100 | 505 | 5.05 |
| SeqAmp CB PCR Buffer | Clontech | 638526 | 1250 µl | 50 µl | 25 | 49 | 1.96 |
| PCR1 and PCR2 primer | Sangon | NA | NA | NA | 200 | 98 | 0.49 |
| Hieff NGS DNA selection Beads | Yeasen | 12601ES56 | 60000 µl | 120 µl | 500 | 943 | 1.89 |
| rRNA and mtRNA sgRNA pool | Sangon | NA | NA | NA | 1000 | 1084 | 1.08 |
| Cas9 nuclease | NEB | M0386T | 500 pmol | 5 pmol | 100 | 195 | 1.95 |
| **DETECTOR-seq** | | | | | | | **21** |

**Table S7. The estimated per sample cost of library preparation for other methods.**

| Methods / Kits | Manufacturer | Catalog # | N sample | $ Total | $ Cost per sample | Reference |
|---|---|---|---|---|---|---|
| SMARTer-seq | Takara | 634413 | 96 | 8642 | 90 | (11) |
| SILVER-seq | NuGEN | 0500-96 | 96 | 13500 | 141 | (10) |
| Phospho-RNA-seq | Illumina | RS-200-0012 | 24 | 6676 | 278 | (9) |

SMARTer-seq, SMARTer Stranded Total RNA-Seq Kit v2-Pico Input Mammalian; SILVER-seq, Small Input Liquid Volume Extracellular RNA Sequencing, whose cost was estimated using Ovation SoLo RNA-Seq Kit; Phospho-RNA-seq, whose cost was estimated using T4 PNK and TruSeq small RNA kit.

**Table S8. Sample and data summary for the development, validation and application of DETECTOR-seq.**

| Stage | Experiments | Figures | Participants (N) | *Plasma* sample N (Pass sample QC) | *EV* sample N (Pass sample QC) | *Plasma* RNA data N (Pass data QC) | *EV* RNA data N (Pass data QC) |
|---|---|---|---|---|---|---|---|
| **1. DETECTOR-seq beta** | RNA extraction comparison | Supplementary Figure 3 | Healthy donor (3) | - | - | 9 | - |
| | CRISPR-Cas9 assisted depletion | Figure 2A | Healthy donor (6) | - | - | 18 | - |
| | DNA digestion with an On-column approach | Supplementary Figure 4 | Healthy donor (7) | - | - | 7 | 4 |
| | | | Colorectal cancer (11) | - | - | 8 | 7 |
| | | | Lung Cancer (13) | - | - | 9 | 7 |
| | | | Total (31) | - | - | 24 | 18 |
| **2. DETECTOR-seq** | *Plasma* versus *EV* analysis with paired samples; Cancer versus Health analysis with all samples | Figure 5, 6, 7 | Healthy donor (31) | 28 | 28 | 19 | 14 |
| | | | Colorectal cancer (24) | 23 | 20 | 23 | 19 |
| | | | Lung Cancer (20) | 20 | 20 | 19 | 19 |
| | | | Total (75) | 71 | 68 | 61 | 52 |
| **3. Benchmarking** | Multiplexing uniformity analysis | Figure 3A | Data generated in stage 2 with DETECTOR-seq | - | - | 6 | 6 |
| | Read retention analysis | Figure 3B | | - | - | 61 | 52 |
| | DNA contamination analysis | Figure 3C | | - | - | 61 | 52 |
| | Accuracy analysis | Figure 3F | | - | - | 61 | 52 |

| | | | | | | | |
|---|---|---|---|---|---|---|---|
| | Consistency analysis | Figure 2D | | - | - | 10 | - |
| | Saturation analysis | Figure 3G, H | | - | - | 12 | 12 |
| | Sensitivity analysis | Figure 3D | Healthy donor (5) | - | - | 25 | - |
| | Reproducibility analysis | Figure 3E | Healthy donor (4) | - | - | 6 | - |

Residual DNA was digested with an On-column approach in DETECTOR-seq beta, while DNA digestion was performed with an In-buffer approach in DETECTOR-seq. EV, extracellular vesicle; QC, quality control.

**Table S9. Clinical characteristics of participants whose *Plasma* or *EV* RNA sequencing data were generated with DETECTOR-seq.**

| | Healthy Donor | Colorectal cancer | Lung cancer |
|---|---|---|---|
| Samples N | 26 | 23 | 20 |
| Age, Mean ± SD | 61 ± 9 | 62 ± 7 | 61 ± 7 |
| % Male | 50% | 52% | 55% |
| % Stage I | - | 22% | 70% |
| % Stage II | - | 26% | 5% |
| % Stage III | - | 39% | 0% |
| % Stage IV | - | 13% | 0% |
| % Stage unknown | - | 0% | 25% |

**Table S10. List of participants whose *Plasma* or *EV* RNA sequencing data were generated with DETECTOR-seq.**

| Sample ID | Group | Age | Sex | Stage | Plasma-SampleQC | EV-SampleQC | Plasma-DataQC | EV-DataQC |
|---|---|---|---|---|---|---|---|---|
| NC-31 | Healthy donor | 59 | M | NA | Pass | Pass | Pass | Pass |
| NC-32 | Healthy donor | 56 | M | NA | Pass | Pass | Pass | Pass |
| NC-33 | Healthy donor | 60 | M | NA | Pass | Pass | Pass | Pass |
| NC-34 | Healthy donor | 62 | M | NA | Pass | Pass | Pass | Pass |
| NC-35 | Healthy donor | 46 | F | NA | Pass | Pass | Fail | Fail |
| NC-36 | Healthy donor | 74 | F | NA | Pass | Pass | Pass | Fail |
| NC-37 | Healthy donor | 66 | F | NA | Pass | Pass | Pass | Fail |
| NC-38 | Healthy donor | 45 | M | NA | Pass | Pass | Pass | Pass |
| NC-39 | Healthy donor | 62 | F | NA | Pass | Pass | Pass | Fail |
| NC-40 | Healthy donor | 71 | F | NA | Pass | Pass | Fail | Fail |
| NC-41 | Healthy donor | 64 | F | NA | Pass | Pass | Pass | Fail |
| NC-42 | Healthy donor | 69 | M | NA | Pass | Pass | Pass | Fail |
| NC-43 | Healthy donor | 67 | F | NA | Pass | Pass | Pass | Fail |
| NC-44 | Healthy donor | 64 | M | NA | Pass | Pass | Pass | Fail |
| NC-45 | Healthy donor | 70 | F | NA | Pass | Pass | Pass | Fail |
| NC-46 | Healthy donor | 52 | M | NA | Pass | Pass | Pass | Fail |
| NC-47 | Healthy donor | 56 | F | NA | Pass | Pass | Pass | Pass |
| NC-48 | Healthy donor | 60 | F | NA | Pass | Pass | Pass | Fail |
| NC-49 | Healthy donor | 58 | F | NA | Pass | Pass | Pass | Fail |

| | | | | | | | | |
|---|---|---|---|---|---|---|---|---|
| NC-50 | Healthy donor | 65 | M | NA | Pass | Pass | Pass | Fail |
| NC-51 | Healthy donor | 70 | M | NA | Pass | Pass | Fail | Pass |
| NC-52 | Healthy donor | 47 | M | NA | Pass | Pass | Fail | Pass |
| NC-53 | Healthy donor | 54 | F | NA | Pass | Pass | Fail | Pass |
| NC-54 | Healthy donor | 73 | M | NA | Pass | Pass | Fail | Pass |
| NC-55 | Healthy donor | 63 | M | NA | Pass | Pass | Fail | Pass |
| NC-56 | Healthy donor | 64 | F | NA | Pass | Pass | Fail | Pass |
| NC-57 | Healthy donor | 41 | F | NA | Pass | Pass | Fail | Pass |
| NC-62 | Healthy donor | 64 | F | NA | Fail | NA | NA | NA |
| NC-63 | Healthy donor | 58 | F | NA | Fail | NA | NA | NA |
| NC-65 | Healthy donor | 51 | F | NA | Fail | NA | NA | NA |
| NC-66 | Healthy donor | 72 | F | NA | Pass | Pass | Pass | Pass |
| CRC-32 | Colorectal cancer | 66 | M | II | Pass | Pass | Pass | Pass |
| CRC-34 | Colorectal cancer | 65 | M | I | Pass | Pass | Pass | Fail |
| CRC-35 | Colorectal cancer | 68 | M | III | Pass | Pass | Pass | Pass |
| CRC-36 | Colorectal cancer | 65 | M | I | Pass | Pass | Pass | Pass |
| CRC-37 | Colorectal cancer | 63 | F | IV | Pass | Pass | Pass | Pass |
| CRC-38 | Colorectal cancer | 56 | M | III | Pass | Pass | Pass | Pass |
| CRC-39 | Colorectal cancer | 63 | M | III | Pass | Pass | Pass | Pass |
| CRC-40 | Colorectal cancer | 41 | F | III | Pass | Pass | Pass | Pass |
| CRC-41 | Colorectal cancer | 49 | F | III | Pass | Pass | Pass | Pass |
| CRC-42 | Colorectal cancer | 64 | F | IV | Pass | Pass | Pass | Pass |

| | | | | | | | | |
|---|---|---|---|---|---|---|---|---|
| CRC-43 | Colorectal cancer | 60 | F | III | Fail | NA | NA | NA |
| CRC-44 | Colorectal cancer | 67 | F | IV | Pass | Pass | Pass | Pass |
| CRC-45 | Colorectal cancer | 64 | M | I | Pass | Pass | Pass | Pass |
| CRC-46 | Colorectal cancer | 68 | F | III | Pass | Pass | Pass | Pass |
| CRC-47 | Colorectal cancer | 65 | M | II | Pass | Pass | Pass | Pass |
| CRC-48 | Colorectal cancer | 61 | M | II | Pass | Pass | Pass | Pass |
| CRC-49 | Colorectal cancer | 69 | F | II | Pass | Pass | Pass | Pass |
| CRC-51 | Colorectal cancer | 64 | M | I | Pass | Pass | Pass | Pass |
| CRC-52 | Colorectal cancer | 52 | F | I | Pass | Pass | Pass | Pass |
| CRC-53 | Colorectal cancer | 62 | F | II | Pass | Pass | Pass | Pass |
| CRC-54 | Colorectal cancer | 56 | F | III | Pass | Pass | Pass | Pass |
| CRC-55 | Colorectal cancer | 62 | M | III | Pass | NA | Pass | NA |
| CRC-56 | Colorectal cancer | 68 | M | III | Pass | NA | Pass | NA |
| CRC-57 | Colorectal cancer | 73 | F | II | Pass | NA | Pass | NA |
| LC-21 | Lung cancer | 60 | M | NA | Pass | Pass | Pass | Pass |
| LC-22 | Lung cancer | 72 | F | I | Pass | Pass | Pass | Pass |
| LC-23 | Lung cancer | 63 | F | I | Pass | Pass | Pass | Pass |
| LC-24 | Lung cancer | 60 | F | NA | Pass | Pass | Pass | Pass |
| LC-25 | Lung cancer | 49 | F | I | Pass | Pass | Pass | Pass |
| LC-26 | Lung cancer | 57 | F | I | Pass | Pass | Pass | Pass |
| LC-27 | Lung cancer | 62 | M | NA | Pass | Pass | Pass | Pass |
| LC-28 | Lung cancer | 54 | F | NA | Pass | Pass | Pass | Pass |

| | | | | | | | | |
|---|---|---|---|---|---|---|---|---|
| LC-29 | Lung cancer | 55 | M | NA | Pass | Pass | Pass | Pass |
| LC-30 | Lung cancer | 59 | F | I | Pass | Pass | Pass | Pass |
| LC-31 | Lung cancer | 74 | F | I | Pass | Pass | Pass | Pass |
| LC-32 | Lung cancer | 73 | M | I | Pass | Pass | Pass | Pass |
| LC-34 | Lung cancer | 60 | M | I | Pass | Pass | Pass | Pass |
| LC-35 | Lung cancer | 53 | M | I | Pass | Pass | Pass | Pass |
| LC-36 | Lung cancer | 57 | M | I | Pass | Pass | Pass | Pass |
| LC-37 | Lung cancer | 52 | M | I | Pass | Pass | Pass | Pass |
| LC-38 | Lung cancer | 60 | F | I | Pass | Pass | Pass | Fail |
| LC-40 | Lung cancer | 67 | M | I | Pass | Pass | Pass | Pass |
| LC-41 | Lung cancer | 67 | M | I | Pass | Pass | Pass | Pass |
| LC-42 | Lung cancer | 73 | M | II | Pass | Pass | Fail | Pass |

QC, quality control; NA, not available.
